# Massive Compression for High Data Rate Macromolecular Crystallography (HDRMX): Impact on Diffraction Data and Subsequent Structural Analysis

**DOI:** 10.1101/2024.09.06.611720

**Authors:** Herbert J. Bernstein, Alexei S. Soares, Kimberly Horvat, Jean Jakoncic

## Abstract

New higher-count-rate, integrating, large area X-ray detectors with framing rates as high as 17,400 images per second are beginning to be available. These will soon be used for specialized MX experiments but will require optimal lossy compression algorithms to enable systems to keep up with data throughput. Some information may be lost. Can we minimize this loss with acceptable impact on structural information? To explore this question, we have considered several approaches: summing short sequences of images, binning to create the effect of larger pixels, use of JPEG-2000 lossy wavelet-based compression, and use of Hcompress, which is a Haar-wavelet-based lossy compression borrowed from astronomy. We also explore the effect of the combination of summing, binning, and Hcompress or JPEG-2000. In each of these last two methods one can specify approximately how much one wants the result to be compressed from the starting file size. These provide particularly effective lossy compressions that retain essential information for structure solution from Bragg reflections.

**Synopsis:** New higher-count-rate, integrating, large area X-ray detectors with framing rates as high as 17,400 images per second are beginning to be available. These will soon be used for specialized MX experiments but will require optimal lossy compression algorithms to enable systems to keep up with data throughput. Some information may be lost. Can we minimize this loss with acceptable impact on structural information?

## 1. Introduction

This paper continues a discussion of approaches to compression of diffraction images in macromolecular crystallography that began with consideration of optimal lossless compression (Bernstein and Jakoncic, 2024). In this paper we consider the increasing need for more lossy compression, arising as data rates increase in macromolecular crystallography and as data retention requirements also increase. Lossy compressions have been in use for a long time, even in crystallography. Reducing a diffraction dataset to a list of structure factors is, indeed, a dramatic lossy compression. With new detectors and many beamline upgrades on the horizon, one will need to use lossy compression to keep up.

The general concept of making “raw” data available for all new PDB structures deposited, as part of improved FAIR^1^ principles may also drive the use of lossy compression. Lossy compression, with the goal of minimal structural information loss, is a viable method to archive diffraction data sets permanently. Here we investigate a few of the most promising available methods to apply lossy compression on two test samples, one used for Sulfur Single-wavelength Anomalous Diffraction (S_SAD) phasing, and one for Molecular-Replacement (MR). We ran tests using various parameters to determine how far we could push the compression ratios and still determine the same structure. It is important to note that, with this approach, the diffraction data that will be stored long-term is exactly the data that was used to solve the structure.

### 1.1. High Data Rate Macromolecular Crystallography

Enabled by changes in technology, macromolecular crystallography increasingly is able to extend its focus from the average state observed in a single crystal, or from merged data from a few crystals, to studies of families of distinct structural states arising from data collection from many crystals. This gradual transition, enabled by brighter sources, smaller beams, and new detectors, is driving a series of disruptive changes in the way diffraction data is collected, processed, and archived. Hardware improvements, such as fast, high-resolution, low-noise and high-dynamic-range detectors, high-brilliance x-ray micro-beams, and automated sample handling combined with autonomous operation, are generating high-data-rate and high-data-volume data streams. Conventional software packages and pipelines, designed for simpler single crystal experiments and one-node serial processing, cannot support or even keep up with such rates and volumes. Networks and computational resources have had to be regularly upgraded. Bottlenecks in data-processing pipelines have been removed, sometimes by converting serial execution to parallel execution on multiple nodes. But those changes themselves have generated yet more network and computational load.

Higher flux, smaller beams, and faster detectors open the door to experiments with very large numbers of very small samples that can reveal polymorphs and time-resolved structural information on dynamic effects, but require re-engineering of approaches to the storing of images. Management of orders-of-magnitude more images and limitations of file systems have favored a transition from simple one-file-per-image systems such as CBF (Bernstein *et al*., 2020) to image-container systems such as HDF5. This further increased the load on computers and networks, and the use of data coming from multiple runs at multiple beamlines has required a re-examination of the presentation of metadata. Over the past few years the high-data-rate macromolecular crystallography community has recognized the importance of complete and consistent metadata. One requires this metadata so datasets can be easily processed at any site from data collected at different times, at another facility, or at multiple facilities, or months and years in the past (Bernstein *et al*., 2020). Full metadata associated with the diffraction data set further makes FAIR principles attainable now.

When we began thinking about this data rates were typically 100 fps and were approaching 1,000 fps in some cases. Now we are facing data rates exceeding 15,000 fps (Hiraki *et al*., 2021). We must either discard a significant portion of the data collected or make serious use of greater compression ratios. Indeed, in both xfel serial crystallography and synchrotron serial crystallography, it has long been acceptable to discard frames that do not appear to contain any significant number of reflections.

#### Terminology of Compression Ratios

For a review of the general terminology of compression see Lelewer and Hirschberg (1987). In this paper we will follow the information-theory approach of Shannon (1948) in which, for this crystallographic application, we focus on the information in diffraction images as streams of numbers representing the intensities of pixels. We define a “compression ratio” as the quotient of the size of the original image to the size of the hopefully smaller compressed image that will be processed or stored. For this to make intuitive sense, one must use comparable units for both the numerator and the divisor, *i.e.* bytes over bytes or bits over bits. For example, for the Dectris Pilatus Detector as released in 2007 (Bernstein, 2010), the original image data as collected consisted of four-byte-long integer pixels, but the data as received had gone through a faithful four-to-one byte-offset compression ratio using the CBF library, i.e. yielding a compression ratio of four-to-one.

Different compression algorithms can achieve different compression ratios for different image data, but if we wish to retain all the information in an image, as Shannon proved, there is an inherent limit to the highest faithful compression ratio that can be achieved, called the “entropy-limit”. Some of the information in the stream may be noise or may be irrelevant to the experiment, and we may choose to “lose” part of the information, giving us a higher compression ratio beyond the entropy limit. In this paper we focus on the extra compression ratio (ECR) achieved beyond the entropy limit. As a practical matter, for most diffraction images, the entropy limit is within a factor of two, or less, of the simple byte-offset compression ratio, so we use the byte-offset compression as our base for computing the ECR. Other compression reference points are possible, but this one is a realistic approximation within a factor of two or less to that with which most macromolecular beamlines currently work.

### 1.2. Lossless Compression

When crystallographic diffraction images are recorded, the common practice has been to retain all details of what has been recorded. Such images have been compressed by lossless compression so that all details are recovered when the images are decompressed. For a general review of lossless compression, see Rahman and Hamada (2019). Bernstein and Jakoncic (2024) discussed how the algorithms related to the lossless Lempel-Ziv-Welch compression algorithm (LZW) (Ziv and Lempel, 1977) (Welch, 1984) are, in many cases, the best choice for a wide range of applications. However, LZW and its derivative algorithms were not implemented for macromolecular crystallography data because of an active patent (Welch, 1985) that would have resulted in significant added costs. As a result, for decades, the MX field employed other algorithms (Abrahams, 1993) (Bernstein *et al*., 1999) (Ellis, Bernstein, 2006). The Dectris Pilatus 6M (Brönnimann *et al*., 2003) (Brönnimann, 2005) adopted the CBF format with byte-offset compression (Bernstein, 2010). By 2012, the LZW patents had expired and available computer hardware could handle higher data-framing rates and more complex compressions. In 2013 Dectris designated Collet’s LZ4 compression (Collet, 2011), a very fast byte-oriented version of Lempel-Ziv compression without an entropy stage, as the compressor for their Eiger detectors (Donath *et al*., 2013). Later the compression ratios were improved by adding a pre-conditioning stage using Masui’s bit-shuffle algorithm (Masui *et al*., 2015). In the interim, others have developed and experimented with further LZW compression algorithms. A popular, fast, and efficient algorithm is Z-standard (“zstd”) (Collet and Kucherawy, 2021).

### 1.3. Lossy Compression

As noted in Kroon-Batenburg and Helliwell (2014), “we have to consider the possibility of lossy compression”. Holton applied a lossy-compression algorithm to ADSC^2^ images from a lysozyme crystal by splitting them into images that contain reflections and those that contain the background^3^. The background images are then compressed so the noise level (*i.e.* the variance of the background) stays roughly the same, and finally the two types of images are recombined. He achieved a compression ratio of 34:1 (original size of data to compressed) without changing the visual perception of the images and without changing the derived structure factors significantly [〈ΔF/σ(F)〉 = 0.6]. Ferrer *et al*. (1998) examined lossy data compression of CCD images with discrete cosine or wavelet transforms, achieving approximately a 30:1 compression ratio. They also found little effect on integrated intensities except for weak reflections, which may be related to altered statistical noise in the background or to the algorithm flattening the weak intensities.

However, the obvious advantages of lossy compression for network transfer and disk space could be outweighed by the (unforeseen) future need for raw images. One consideration could be the diffuse scattering of less-ordered crystals that occurs in areas that we normally designate background and which is weak and varies slowly. Currently, no one has established the extent to which such compression algorithms will significantly affect diffuse intensities in reciprocal space. Underwood *et al*. (2023) have proposed combining lossless compression in rectangles around Bragg peaks with lossy compression for the remaining background to cope with the coming data volume loads for serial crystallography. They call their algorithm ROIBIN-SZ. Galchenkova and co-workers (2024) considered use of lossy compression in serial crystallography and saw encouraging results with “real time hit finding, binning, quantization to photons, and non-linear reduction of the dynamical range,” but noted a significant loss of resolution achieved with ROIBIN-SZ in comparison to techniques that preserve the background.

Fields such as astronomy that must deal with very large volumes of mostly empty image data have long employed lossy compression; a popular example of useful software is Hcompress (White *et al*., 1992). Hcompress is a method which is based on the Haar - wavelet-based Haar transformation (Fritze *et al*., 1977). The Haar wavelet is a flat-topped wavelet with discontinuous transitions that help in modelling sharp peaks such as stars and potentially diffraction-peak images. The purpose of Hcompress was to compress astronomical images quickly, reducing the redundancy of pixels displaying bright celestial object images. In addition to that, astronomical images contain a lot of noise in the dark-sky background, which can be removed. These characteristics of astronomical images are shared by macromolecular crystallographic diffraction images.

As we will see, our experiments bear out the value this suggests in using Hcompress for MX. Applying these approaches can result in significant cost savings. See the section on **Worked Example of Estimated Costs at AMX** In the supporting information. If we assume a data-management policy requiring very long term storage of data, but that the data can move to lower-cost off-line storage a decade after collection, the current annual cost estimate for archival lossless storage at USD 14,400 per annum per beamline can be converted to a steady state cost of USD 144K per annum per beamline. This could be reduced to less than USD 1000 per annum per beamline if lossy compressions with ECRs averaging more than 144:1 prove satisfactory.

For all examined lossy compression schemes, the computational resources devoted to data compression are small compared existing needs such as data reduction and image scoring. Furthermore, these computational resources can be deployed during periods when data acquisition is not occurring, such as scheduled equipment maintenance periods. Hence, we believe that the computational overhead needed for compression can be negligible.

Decompression will be built into data processing plugins, essentially resulting in no decompression penalty for all users.

### 1.4. The FAIR Principle and the United States White House Mandate

There are two factors in tension with any data-management approach that discards any diffraction images or even any portions of diffraction images. There has been a world-wide movement of science towards the so-called “FAIR” principles (Wilkinson *et al*., 2016), which require that scientific data be Findable, Accessible, Interoperable, and Reusable. In 2022, the United States formally joined this movement with the White House mandate that data from Federally funded research would become freely available by the end of 2025 (White House, 2022). Clearly, we must devise some rational structure to achieve an appropriate balance between the desire to share all data and the availability of the necessary resources.

Making the right choices in lossy compression should help to achieve that appropriate balance. For MX there are some special issues to consider in achieving the necessary balance. In some protein dynamics studies one needs to keep all diffraction data from all samples with sufficient data to show the aspects of dynamics supported by non-Bragg scattering. One must also reveal aspects supported by careful clustering of families of multiple states, shown in appropriately chosen Bragg diffraction images for which most of the background pixels can be discarded. In addition, for non-dynamic studies, most images do not provide useful data, and most pixels in the images that do provide useful data are not relevant to the final results. It should be sufficient in the course of applying lossy compression to retain an appropriately chosen subset of between one-half and one percent of the images as lossless compression images, in order to detect the cases in which non-Bragg background data is relevant and scientifically interesting.

Here, we focus our attention on a few types of lossy compression: frame summing, pixel binning, JPEG-2000, and Hcompress, and optimal combinations of these. We report the effects of these compressions on analysis and structural refinement for two complete representative test data sets. We have found a few promising lossy compressions that can be used routinely, depending on the type of experiment planned or on the intensities of the observed reflections.

Future steps are to investigate the impact of lossy compression choices on real-life data collections by making lossy compression options available, first as redundant parallel paths in data-processing production pipelines, and then, if justified by the results, as selectable alternative processing choices; this builds on Holton’s approach of segregating the background processing from reflection processing.

## 2. Materials and Methods

### 2.1. Data Collection

Data were collected at the National Synchrotron Light Source II (NSLS-II) Highly Automated Macromolecular Crystallography (AMX) beamline 17-ID-1 (Schneider *et al*., 2022) at Brookhaven National Laboratory, using an Eiger X 9M detector (Förster *et al*., 2016). We collected two distinct data sets: 1) a data set from an HIV reverse transcriptase crystal (Hollander *et al*., 2023) (Prucha *et al*., 2023) (Chan *et al*., 2020) (Duong *et al*., 2020) at 13.5 keV for molecular replacement studies, and 2) a redundant data set on a lysozyme crystal at 7.5 keV to increase the anomalous scattering factor of Sulfur for Sulfur SAD phasing. There were a total of 1200 and 1800 0.2-degree of rotation frames each for the HIV reverse transcriptase and the low energy S_SAD lysozyme samples, respectively.

### 2.2. Lossy Compression Methods Tested

#### 2.2.1. Image summing

When data have been collected using fine χπ-slicing (0.2 degrees / frame), a very effective compression is to sum a small number of sequential frames, pixel-by-pixel. For this series of experiments, we achieved this summing with the program merge2cbf from the XDS package [https://xds.mr.mpg.de/html_doc/merge2cbf_program.html]. For the HIV reverse transcriptase we tried summing by 2’s, by 5’s, and by 10’s, and experimented with summing by 20’s and 40’s, essentially converting the angle of each slice to an angle two, five, or ten times as coarse, as well as even coarser slices (20 and 40). Recall that in the 1980s, employing x-ray film as detector, we might have taken one-to-five-degree images^4^, the equivalent of 5 – 25 of these small frames. We label n-fold coarse slicing “SUMn”. For moderate values of n, this has very little effect on the quality of the peaks collected other than to make it less likely to encounter “partial reflections,” but it does cause a linear increase in the level of the background, making it more challenging to detect weak peaks.

#### 2.2.2. Pixel binning

Another very effective compression is to sum pixels in the plane of the image which we call “binning”. For example, in 2x2-fold binning, each even numbered pixel in each even numbered row is replaced by the sum of that pixel with the pixel to the right and the pixel above and the pixel diagonally above and to the right. We label that 2-fold binning “BIN2”, and in general label n-fold binning “BINn”. This has the effect of reducing the number of pixels by a factor of four, or equivalently of increasing the size of each pixel by a factor of two in each direction. We experimented with BIN2, BIN4, BIN6, BIN8, and BIN10 for the reverse transcriptase. For the lysozyme data set, we only applied BIN2.

By combining the coarsening of the slicing with binning by two, one can achieve a compression by a factor of eight (SUM2 + BIN2) or twenty (SUM5 + BIN2) at very low cost in terms of both time and statistics. For these experiments the binning was done by use of the program cif2cbf from the CBFlib package [https://github.com/yayahjb/cbflib]. The use of modest amounts of coarsening of the slicing with binning combines very well will any of the popular lossless compressions including those from CBFlib as well as with bslz4 and bszstd.

#### 2.2.3. JPEG-2000 and Hcompress

Two compression techniques from other scientific fields may be useful for diffraction images: JPEG-2000 from the image-processing industry (Santa-Cruz *et al*., 2002) as released in the OpenJPEG package (ISPGroup, 2023), and Hcompress from astronomy (White *et al*., 1999), (White, 2019). JPEG-2000 uses the Daubechies wavelet, which is a smoother generalization of the Haar wavelet, which is now referred to as DB-1.

For these studies, JPEG-2000 requires a preliminary conversion of images to 16-bit grey scale tiff format (Data, 1947), while Hcompress requires a preliminary conversion to astronomical fits format (Pence *et al*., 2020) after conversion to tiff. We used the siril package for conversion from tiff to fits [https://siril.org/]. (For our data, use of 8-bit grey scale tiff proved to be too aggressive a lossy compression compared to 16-bit tiff, though it might be suitable for other data with less dynamic range and a flatter background.)

For both samples, we tried various compression levels available in the JPEG-2000 and Hcompress applications.

Finally, to study higher compression ratios, we tried various combinations of summing, binning, JPEG-2000, and Hcompress. In these the combinations of summing pairs of images, 2x2 binning and Hcompress were particularly interesting in terms of high compression ratios with good data processing statistics. We label JPEG-2000 compression with a target compression ratio of n-to-1 as “J2Kn”. We label Hcompress compression with m-fold scaling as “HCOMPm”.

We tried various combinations of compressions on a lysozyme dataset of 1800 frames and on an HIV reverse transcriptase dataset of 1200 frames, both collected in 0.2-degree per frame, for 360 degrees of lysozyme and 240 degrees of HIV reverse transcriptase. Examples of uncompressed native lysozyme, BIN2_SUM2 (eight-to-one compressed) lysozyme, uncompressed native HIV reverse transcriptase, and BIN2_SUM2 (eight-to-one compressed) HIV reverse transcriptase are shown in Figs 1-a, 1-b, 1-c and 1-d, respectively. Note that in each case the compressed images preserve almost all significant reflections against backgrounds stronger than in the native images. Notice as well that the two native images differ significantly in terms of density and intensity of peaks across the images and uniformity of the backgrounds.

**Figure 1.**
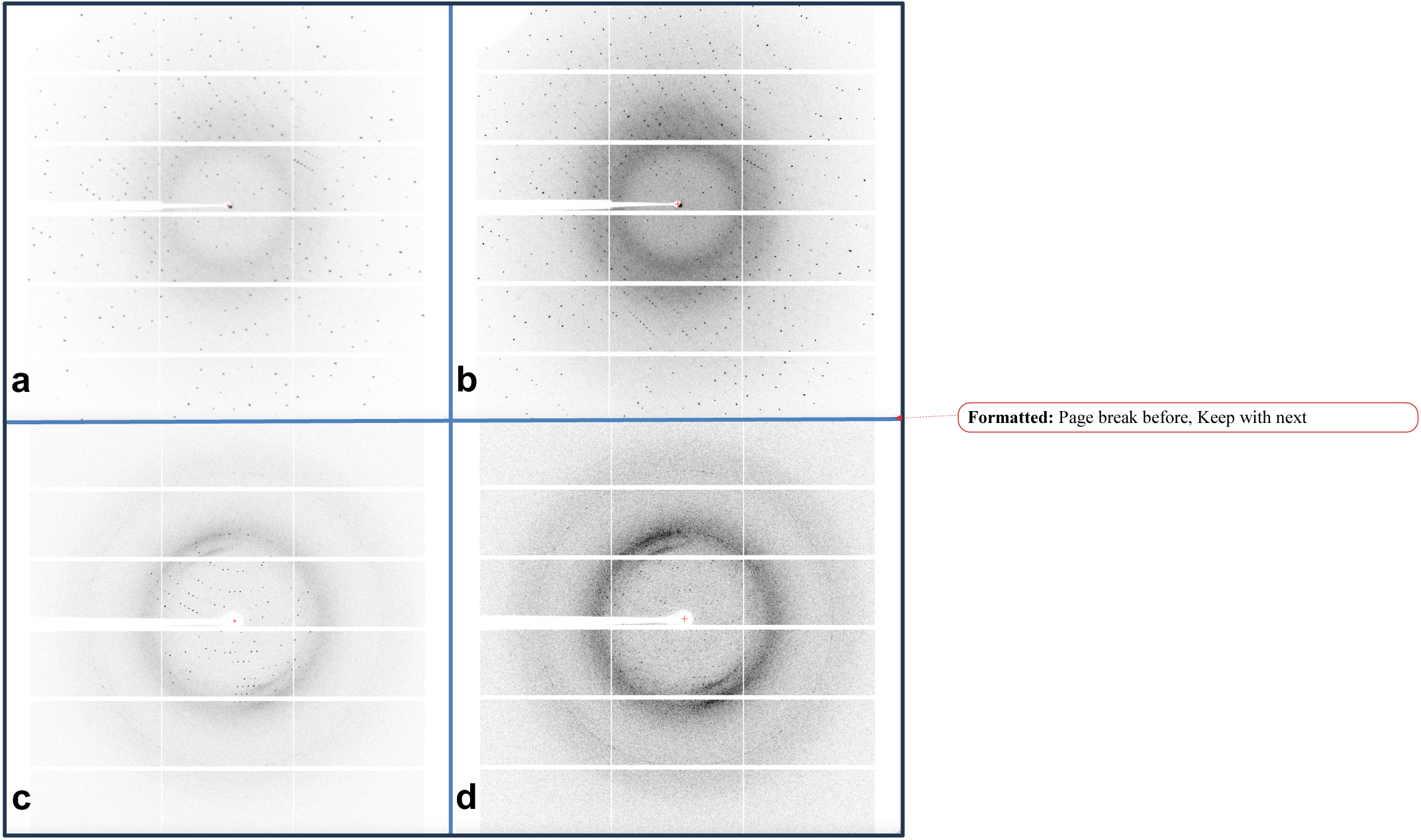
a. Lysozyme frame 900 of 1800 from the native dataset. b. Lysozyme frame 450 of 900 from the 8:1 compressed BIN2_SUM2 dataset; this is the Fig 1 image, added to the one after it, and then with pixels summed 2x2. Notice the much stronger background with well-preserved peaks over the entire image, and that one can pick out a few more peaks in b than in a. c. HIV reverse transcriptase frame 600 of 1200 from the native dataset. d. HIV reverse transcriptase frame 300 of 600 from the 8:1 compressed BIN2_SUM2 dataset. Notice the much stronger background in d than in c.

All 1800 original diffraction frames for the S_SAD lysozyme, are available on zenodo^5^ with all compressions applied and the data back-converted to CBF format.

### 2.3. Data Processing and Analysis

The two original data sets (hdf5 files) were analyzed with the fast_dp pipeline (Winter and McAuley, 2011). An XDS.INP file was derived from the intermediate XDS.INP files that fast_dp creates for spot finding, indexing and integration. Execution of XDS generated XDS_ASCII.HKL files which contained corrected intensities. These were processed further using pointless and aimless for space-group selection and for scaling. We then used the two resulting MTZ files, one for the reverse transcriptase and one for the lysozyme, for structure solution.

The reverse transcriptase structure was refined using PDB (Bernstein *et al*., 1977) (Berman *et al*., 2003) entry 8STS (Hollander *et al*., 2023) as a starting model for the molecular refinement. The refined model was manually corrected to account for a loop/domain motion using COOT (Emsley and Cowtan, 2004). The final model was refined using REFMAC (Murshudov *et al*., 1997) (Vagin *et al*., 2004) and COOT.

The lysozyme uncompressed data were processed using the same manual workflow consisting of XDS, pointless, and aimless. The reduced file was used for a SAD-phasing experiment using the hkl2map (Pape and Schneider, 2004) and the SHELXC, SHELXD, and SHELXE components of the SHELX system (Sheldrick, 2008) software packages. We selected these packages since they offer manual tuning of some parameters. The output file was used for further processing using density modification (DM) (Cowtan, 2010) and model building from ArpWarp (Langer *et al*., 2008) using a semi-automated workflow.

All lysozyme and HIV reverse transcriptase compressed data were reduced using the corresponding XDS.INP file with a few modifications to account for binning and summing when required. Pixel sizes, number of pixels, beam center, and detector gaps, plus oscillation range and number of frames were updated accordingly. No additional modifications to XDS.INP were required for the JPEG-2000 and HCOMP compression *per se*.

Subsequent models from the reverse transcriptase compressed data were refined using the original model as a starting point for rigid-body and atomic refinement using REFMAC. All lysozyme compressed reduced data were used following the same workflow employed to solve the uncompressed S_SAD.

## 3. Results

### 3.1. Achieved Compression Ratios

As one might expect from the differences in the native images in both datasets (Figs. 1.a. and 1.c), the extra compression ratios achieved from the two datasets, as shown in Figs. 2 and 3, the lysozyme dataset is a little more difficult to compress that the HIV reverse transcriptase dataset. Many of the available extra compression ratios tested exceeded 500:1. An appropriate subset of these compressions were tested for their impact on the effectiveness and quality of structure solution.

**Figure 2.**
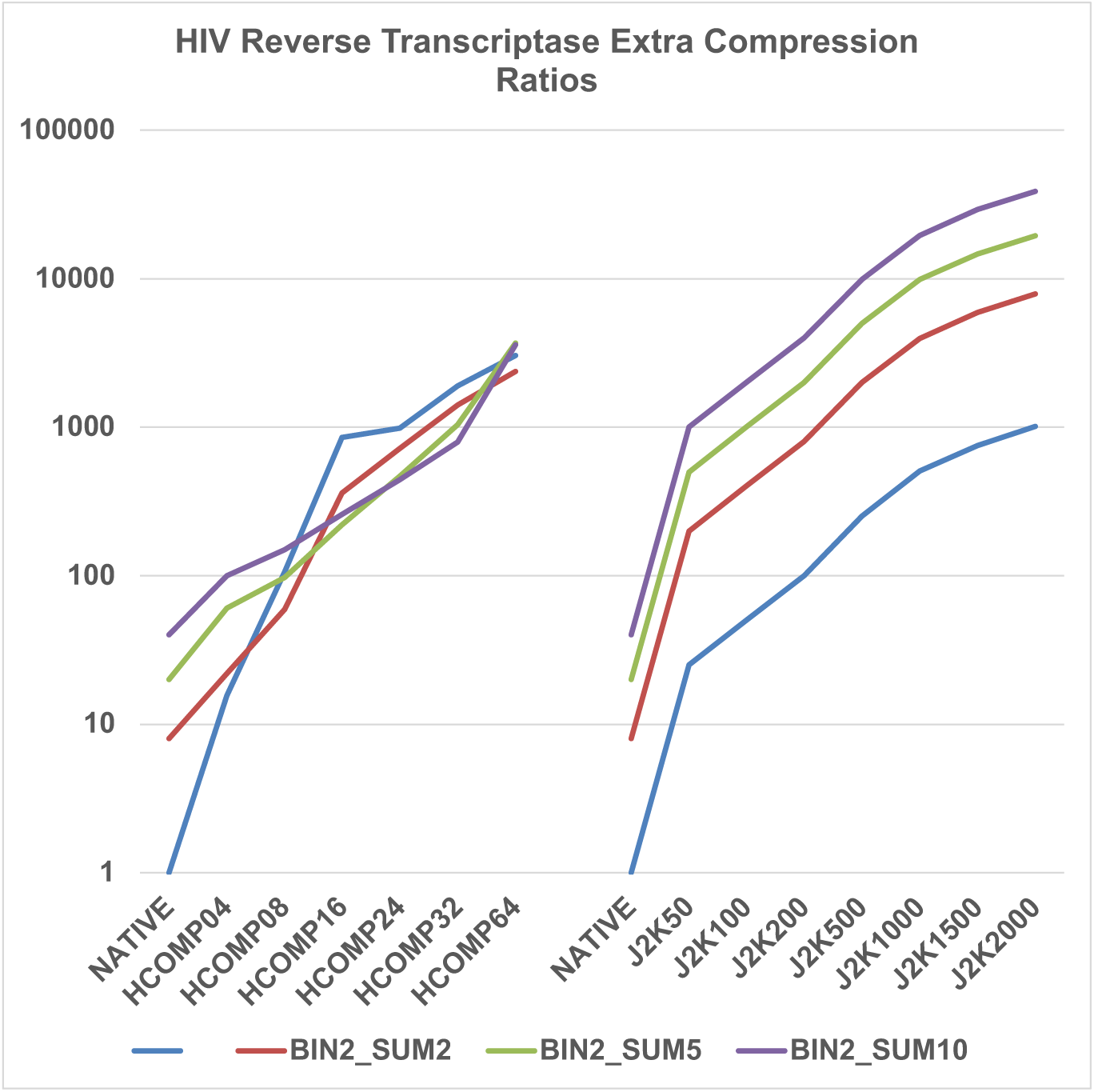
Wide selection of available lossy compressions for HIV reverse transcriptase. Note that many of the compression ratios exceed 500:1. Not all compressions tested are included in this Fig. A Table with all compressions is included in the supplemental material.

**Figure 3.**
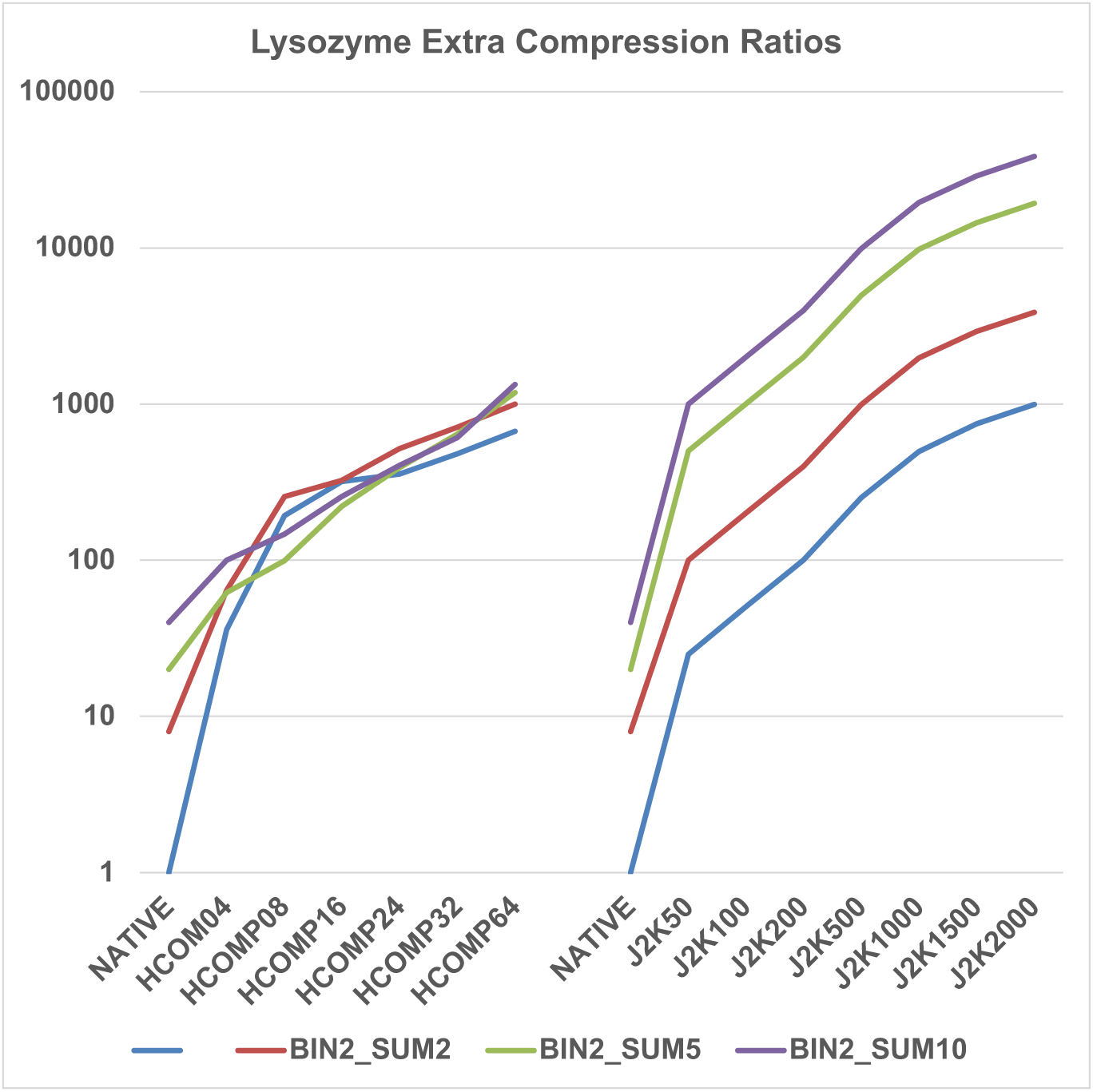
Wide selection of available lossy compressions for lysozyme. Note that many of the compression ratios exceed 500:1. Not all compressions tested are included in this figure.

We are surveying the impact of lossy compressions on data reduction as well as on the ability to solve 3-D crystal structures. One can gauge lossy compression effects in several ways: 1) first looking at the corresponding overall diffraction pattern, 2) then observing impacts on weak and strong reflections, observing the diffraction image of the integrated reflection intensity divided by surrounding background, by looking at data reduction key parameters, and 3) finally looking at the refined structure analysis and the corresponding electron density maps.

### 3.2. Compression Results for Lysozyme

The native and compressed data for lysozyme were processed with XDS (Kabsch, 2010), aimless, pointless, SHELXC, SHELXD, and SHELXE components of the SHELX system (Sheldrick, 2008), and other CCP4 components (Evans, 2014).

Fig. 4 shows the data reduction statistics and extra compression ratios for lysozyme data sets using various compression methods. The corresponding Tables S4 and S5 in the supplementary material highlight the most commonly used parameters (Rpim (precision indicating merging R), completeness, mean I/σ(I), unique reflections) as well as a subsequent set of parameters to illustrate potential damage to data statistics prior to S_SAD phasing (anomalous correlation and mid slope anomalous probability). The objective is to get the maximum compression that is consistent with keeping Rpim low and both I/σ(I) and

**Figure 4.**
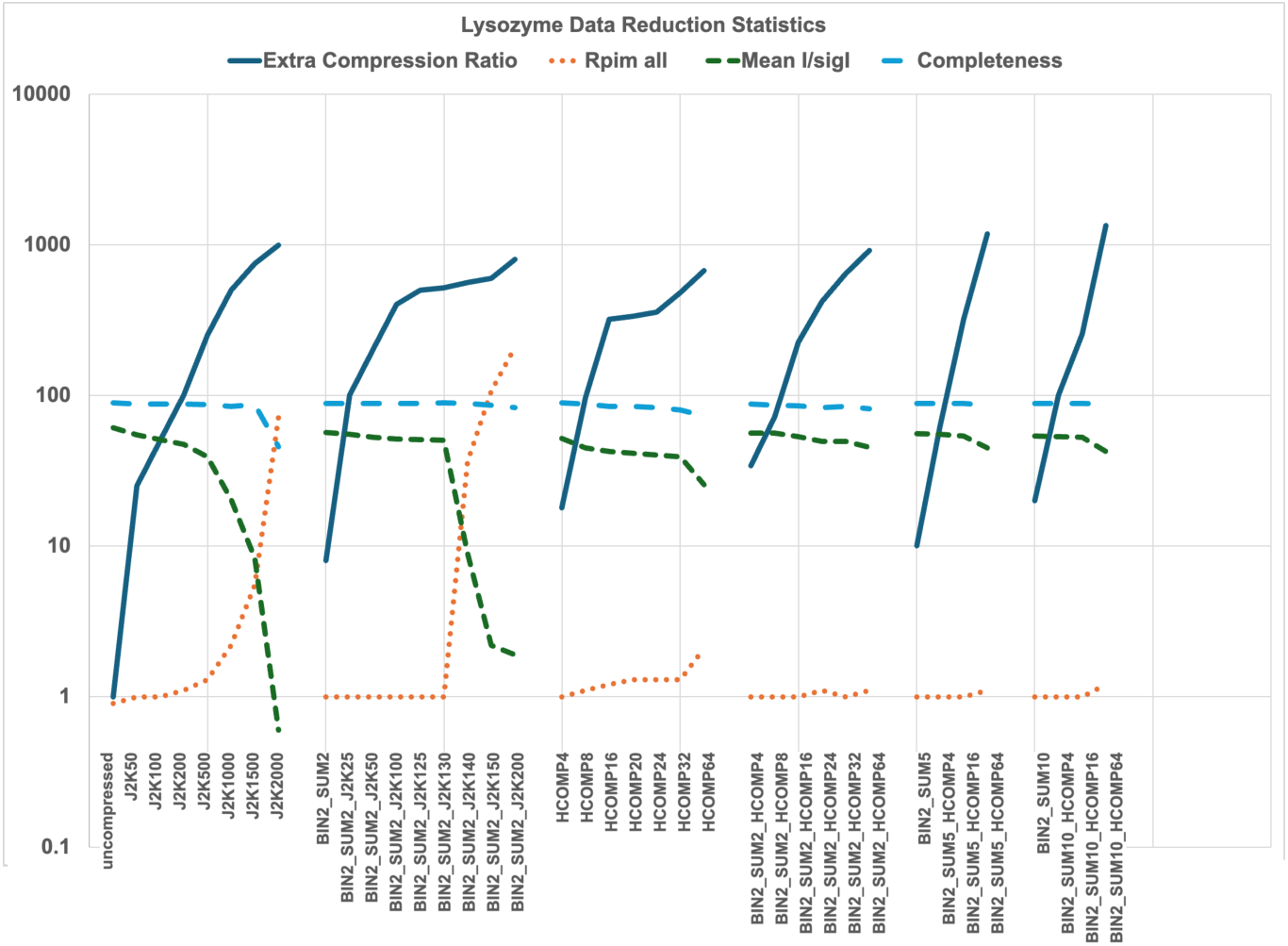
Shows a portion of the data reduction statistics for all lysozyme data sets. The figure shows the most commonly used parameters, Rpim, completeness, mean I/σ(I) and completeness. The vertical axis is the numeric value for the extra compression ratio achieved as well as for each of the commonly used parameters. More complete information is given in the supplementary material.

Completeness high. Numbers for the highest resolution shell (1.67 – 1.64 Å) are shown there in parentheses. For simplification, other parameters usually displayed are not included in Tables S4 and S5. 1800 frames were collected at 7.5 keV to improve Sulfur dF” to maximize likelihood of successful phasing using anomalous signal from the 10 S atoms present in the lysozyme. Note that the protein was crystallized in a NaCl solution resulting in the observation of seven additional peaks.

Here we are investigating the effects of binning pixels, summing frames, JPEG-2000, and Hcompress lossy compressions, and subsets of combinations. The AMX beamline relies on an EIGER X 9M detector with a minimum distance of 100 mm between the sample and the detector, limiting the highest achievable resolution for a given wavelength. This results in overall completeness of 88.9 % in the 20.00 – 1.64 Å range.

Note that after we first attempted lossy compression on the HIV reverse transcriptase, we concluded that some of the compressions tested (BIN4, BIN6, and higher; various summing sets) were not likely to be useful in the lysozyme case. Therefore, for lysozyme we focused on Hcompress, JPEG-2000 and the combinatorial effects of BIN2_SUM2, BIN2_SUM5, BIN2_SUM10 with either JPEG-2000 or Hcompress.

Fig. 4 and Tables S4 and S5 highlight the most important parameters selected to probe effects of lossy compression when compared with the “uncompressed” data. From the data reduction statistics perspective only, it appears that all compressions tested have effects on the data statistics, varying effects that do increase with increased level of compression from JPEG-2000 and Hcompress.

#### 3.2.1. JPEG compression

For lysozyme, we tested JPEG-2000 compression levels ranging from 50:1 to 2000:1; measured compression ratios were consistent with the ratio requests, i.e. the extra compression ratios were approximately half the requested J2K overall compression ratios. Applying increasing JPEG-2000 compression levels resulted in gradual degradation of I/σ(I) and R factors, with J2K2000 data being severely damaged; one might even say destroyed. Completeness of J2K2000 data was half that of the uncompressed data, and R factors as well as I/σ(I) were essentially those of random intensities. Focusing on the highest resolution shell, any level of JPEG-2000 compression generated measurable error. That said, we are interested in the measurable impact on the structural information that one can derive from the compressed data and comparing it with the uncompressed data. This will inform us as to safe levels to use in future experiments.

Low levels of compression (50, 100, and 200) appear to alter data statistics only marginally. Most crystallographers would still find the data “good” according to all the metrics included in Fig. 4 and Tables S4 and S5.

The “interesting” area is with moderate JPEG-2000 requested compression ratios, 500, 1000 and 1500. J2K500 (∼250 to 1 extra compression ratio) statistics are superficially sufficient to allow for structure solution. However I/σ(I) was significantly reduced, from 61 to 39. For J2K1500 (∼750 to 1 extra compression ratio) the I/σ(I) value is is reduced drastically to 28.3 and reflection intensities and surrounding background levels were significantly impacted.

Nevertheless, we attempted to solve the S_SAD lysozyme structure using all generated compressed data. Some of the over-compressed cases failed. The less-compressed, successful cases are bolded in the table in the supplementary data.

#### 3.2.2. Hcompress compression

For Hcompress the amount of compression is specified as “scales”, rather than target compression ratios. Hcompress compression scales ranging from 4 to 64 were applied to the uncompressed data, corresponding to compression ratios ranging from 36 to 1344. Unlike JPEG-2000, which did not impact the overall completeness, Hcompress on the other hand results in gradual loss of completeness with increasing scale of compression. Further, R factors are moderately impacted by the higher compression scales.

These two algorithms clearly display different effects on diffraction data that have been investigated at the various Hcompress reflection scales. See Discussion Section 4.

Learning from the compressions that were first tested for the HIV reverse transcriptase we limited compression algorithms and combinations to a selected set. Specifically, summing frames (2, 5, 10, 20 and 40) and binning pixels (2,4 and 6) were not tested for lysozyme. We focused on combinations of 2x2 binning and summing by 2 without and with various Hcompress scales and JPEG-2000 target compression ratios. Higher levels of summing in combination with 2x2 binning and Hcompress were tested to evaluate impact of greater lossy compression for S_SAD phasing.

#### 3.2.3. Binning / Summing / JPEG 2000

We applied JPEG-2000 compression to the “BIN2_SUM2” frames with JPEG-2000 target compression ratios ranging from 25 to 200. Data statistics included in Fig. 4 and Tables S4 and S5 compared to the BIN2_SUM2 the uncompressed data remained “unperturbed” until we applied J2K140 or higher. Starting at target compression ratio 140, R factors, and I/σ(I) represent statistics from random diffraction with decreasing quality for increasing compression level. It appears that the transition from level 130 to 140, for the lysozyme 7.5 keV dataset for S_SAD phasing, corresponds to an inflection point. This transition will be investigated further in terms of Bragg reflections intensities and local background profile. These correspond to compression ratios of 1039 (level 130) and 1119 (level 140). Discussion of the effect of higher levels of compression from either JPEG-2000 or Hcompress is included below.

#### 3.2.4. Binning / Summing / Hcompress

As for the JPEG-2000 combined case, we tested six Hcompress scales, ranging from 4 to 64. Like the JPEG-2000 case, most compressed data processed with “unnoticeable” damage when focusing on the parameters in Fig. 4 and Tables S4 and S5. We observe gradual noticeable decay of the high resolution statistics as well as the overall completeness and I/σ(I) but with no measurable impact on experimental phasing and subsequent automated model building. Compression ratios ranging from 34 to 915 were achieved. Unlike for JPEG-2000, for Hcompress we also investigated the following combinations: BIN2_SUM5 and BIN2_SUM10 with Hcompress levels of 4, 16 and 64. These correspond to compression ratios ranging from 62 to 1336.

### 3.3. Lysozyme Lossy Compression S_SAD Phasing

Starting with a S_SAD lysozyme structure, we investigate the effect of lossy compression of this data set on the solution of the S_SAD structure using a combination of four compression algorithms with compression ratios ranging from 50 to 2672. Two main algorithms were tested, JPEG-2000 and Hcompress, each with different impact. Not counting the native CBF dataset all 37 processed data sets were subsequently used for phasing experiments following the workflow derived from the uncompressed data.

As shown in Table 1 and supplementary Tables S1, S2, and S3, the uncompressed data produced 17 correct SHELXD heavy atom searches. 37 lossy compressions were tested, with extra compression ratios ranging from 25 to 1336 and a subset of these are included in Table 1. A large number of the lossy compressions produced equally good heavy atom search solutions, with compression ratios up to almost 500:1. See Tables S1, S2, and S3 in the supplemental information for additional compression runs, many of which were successful and some of which failed.

**Table 1.** A sampling of the successful processing of lysozyme compressed data sets, following the standard workflow: XDS, pa_sg_res_cut -S P4_3_2_1_2, mtz2sca, aimless.mtz, aimless.sca, and using hkl2map for SHELX CDE. Note that pa_sq_res_cut^6^ is a bash shell script wrapper for aimless and pointless, for the downstream processing of fast_dp output. This script allows for trimming integrated reflection files (from XDS) based on resolution and frame number, in addition to space group assignment. The number of heavy atom solutions from SHELXD (with CC weak greater than 15) is given, along with the number of residues traced by SHELXE, connectivity of map, dm.mtz FOM, number of residues autobuilt using ArpWarp, and the final Arp model R-value. See Tables S1, S2, and S3 in the supplemental information for additional compression runs, many of which were successful and some of which failed.

| Compression | Extra Compression Ratio | SHELXD Sol'ns | SHELXE Resid. | SHELXE FOM /CONN | DM FOM | ARP res /correct | R |
| --- | --- | --- | --- | --- | --- | --- | --- |
| Uncompressed | 1 | 17 | 119 | 0.65/0.77 | 0.83 | 127/1.00 | 22.4 |
| BIN2_SUM2_HCOMP04 | 17 | 2 | 106 | 0.65/0.76 | 0.83 | 127/1.00 | 23.5 |
| BIN2_SUM2_HCOMP08 | 36 | 11 | 109 | 0.60/0.76 | 0.82 | 127/1.00 | 23.6 |
| BIN2_SUM2_HCOMP16 | 112 | 15 | 103 | 0.63/0.76 | 0.82 | 127/1.00 | 23.1 |
| BIN2_SUM2_HCOMP24 | 211 | 14 | 99 | 0.61/0.76 | 0.82 | 128/0.999 | 26.7 |
| BIN2_SUM2_HCOMP32 | 323 | 10 | 112 | 0.65/0.76 | 0.81 | 127/1.00 | 23.7 |
| BIN2_SUM2_HCOMP64 | 458 | 1 | 105 | 0.63/0.76 | 0.79 | 127/1.00 | 24.0 |

Two additional parameters, related to potential impact on phasing experiments were included in Tables S4 and S5; the correlation of the intensities of two randomly selected half data sets between Friedel-mates (CC½ of anomalous data) and the mid-slope of the normal probability of anomalous difference, both provided by aimless.

#### 3.3.1. Uncompressed data Sulfur_SAD phasing and automated model building

The heavy atom search was performed searching for 14 S atoms, using 1000 trials unless otherwise specified. Phasing, density modification (DM) and chain auto-tracing were performed with 200 cycles of density modification with 40 % solvent fraction with three cycles of tracing. DM was executed using the output from SHELXE for 200 cycles using 40 % solvent fraction and subsequently used for model building in ArpWarp using default settings. The lysozyme data set was acquired at 7.5 keV to maximize anomalous contribution from the 10 S atoms and the various Cl ions that are often observed when crystals are obtained from NaCl precipitant based solution. In SHELXD, 17 out of the 1000 trials were successful sub-structure solutions with the best one saved and used in SHELXE. 119 out of 129 amino acids were traced in the density which displayed a connectivity of 0.77 and a phase FOM of 0.65. ArpWarp placed and refined 127 amino acids including side chains with a crystallographic R factor of 22.4. In other words, the uncompressed data resulted in straightforward experimental phasing and automated model building.

#### 3.3.2. Compressed data Sulfur_SAD phasing and model building

Phasing was attempted with all compressions tested and phasing results (pass or fail; pass defined as successful experimental phasing including heavy atom search, phasing and chain tracing) were used to perform further higher levels of compression and explore additional combinations not available for the HIV reverse transcriptase. Here, it is important to distinguish two important steps relevant to successful phasing experiments: sub-structure solution followed by phasing and density modification. Of the 37 compressions tested (with ratios ranging from 8 to 1336), there were 27 successful runs that resulted in straight forward sub-structure solutions followed by essentially indistinguishable phasing results compared to the uncompressed data. These 27 compressions with “insignificant” structural changes had ratios ranging from 8 to 1336. It is key to emphasize that summing frames combined with binning pixels in those frames does significantly increase the intensities of all reflections with the most impact on partial strong reflections recorded over a few consecutive frames. When summing and binning were combined with Hcompress or JPEG-2000, higher compression ratios were achieved with minimum impact on phasing results.

JPEG-2000 with target compression ratio greater than 1500, Hcompress with a scale of 64 as well as BIN2_SUM2 combined with JPEG-2000 J2K*xxx* with *xxx* of 140 or greater did not yield successful sub-structure solution and phasing. Attempts to use up to 10,000 SHELXD trials and good sub-structure solution failed.

A few compressions tested required up to 10,000 SHELXD trials for a successful substructure solution needed for successful phasing and model building. These compressions were: HCOMP16 and HCOMP20 scales, BIN2_SUM5 with HCOMP4, as well as BIN2_SUM10 with and without HCOMP4.

The remainder of the compressions tested failed to yield sub-structure solutions. However, when using a set of known heavy atom positions from the previous SHELXD step, we were able to use SHELXE for phasing and phase improvement. In particular as shown in supporting information tables S1 and S2, in the blue background areas, solution of the lysozyme data after lossy compression with JPEG-2000 at target compression ratios of 500:1, J2K500, and 1000:1, J2K1000 (actual extra compression ratios of 251:1 and 498:1 respectively) needed the heavy atom positions from the solution to the JPEG-2000 200:1 target compression ratio, J2K200, compressed data (actual extra compression ratio 100:1), and solution of the lysozyme data after lossy compression with Hcompress at scales of 24, HCOMP24, and 32, HCOMP32 (actual extra compression ratios of 357:1 and 480:1 respectively) needed the heavy atom positions from the solution to the Hcompress scale 8, HCOMP08, compressed data (actual extra compression ratio 96:1).

Essentially, very high level of compressions impact diffraction peaks and background so that the data is unsuitable as observed during reduction and subsequent analysis for use in experimental phasing. The unsuitable cases were JPEG-2000 extra compression of 1000:1 with achieved extra compression ratio of 500:1, HCOMP64 achieving compression ratio of 336, and BIN2_SUM2_J2K*xxx* with *xxx* than 140 corresponding to extra compression ratios greater than 225.

BIN2_SUM2 images generate stronger peaks than uncompressed data. Therefore BIN2_SUM2_J2Kxxx for higher values of xxx tends to impact counts on the shoulders of the reflections rather than the reflections themselves.

HCOMP64 significantly reduces the completeness of the data when applied to uncompressed data, but HCOMP64 when applied after BINx_SUMx does not significantly impact completeness.

***That is, the achieved extra compression ratio is not, by itself, a good metric to determine whether or not phasing experiments can then be carried out, it is important to consider what compression algorithms are being used in what order and what they do to the reflections. See the discussion of reflections, below***.

### 3.4. Compression Results for HIV Reverse Transcriptase

The HIV reverse transcriptase dataset presents a molecular replacement problem for phasing. Because we were looking for a “canary in a coal mine” effect, we were unprepared for the resilience with which modern data reduction and structure solving software were able to handle highly compressed data. Our first benchmarking trial used total compression ratios only up to 200 (extra compression ratio 100), which proved insufficient to unambiguously probe the maximum limits of acceptable compression for two of our compression strategies (JPEG-2000 and HCOMPRESS). We then repeated our benchmarking to compression ratio up to 500 (extra compression ratio 250). However, by combining multiple different compression strategies in sequence, we found that in some cases even 500 was insufficient to detect the onset of detectable degradation. Hence, we repeated the entire exercise again with a maximum compression ratio for JPEG-2000 of 2000 (extra compression ratio 1000). Use of BIN2_SUM2 allowed us to reach higher extra compression ratios by a factor of as much as eight. See Figs. 6 and 7. **Although we did not exhaustively test all possible combinations of compression strategies, we believe that the combination of two-fold data binning followed by two-fold frame binning followed by HCOMPRESS is the optimal strategy for lossy compression, which maximizes the total compression ratio with the least degradation of data quality.**

**Figure 5.**
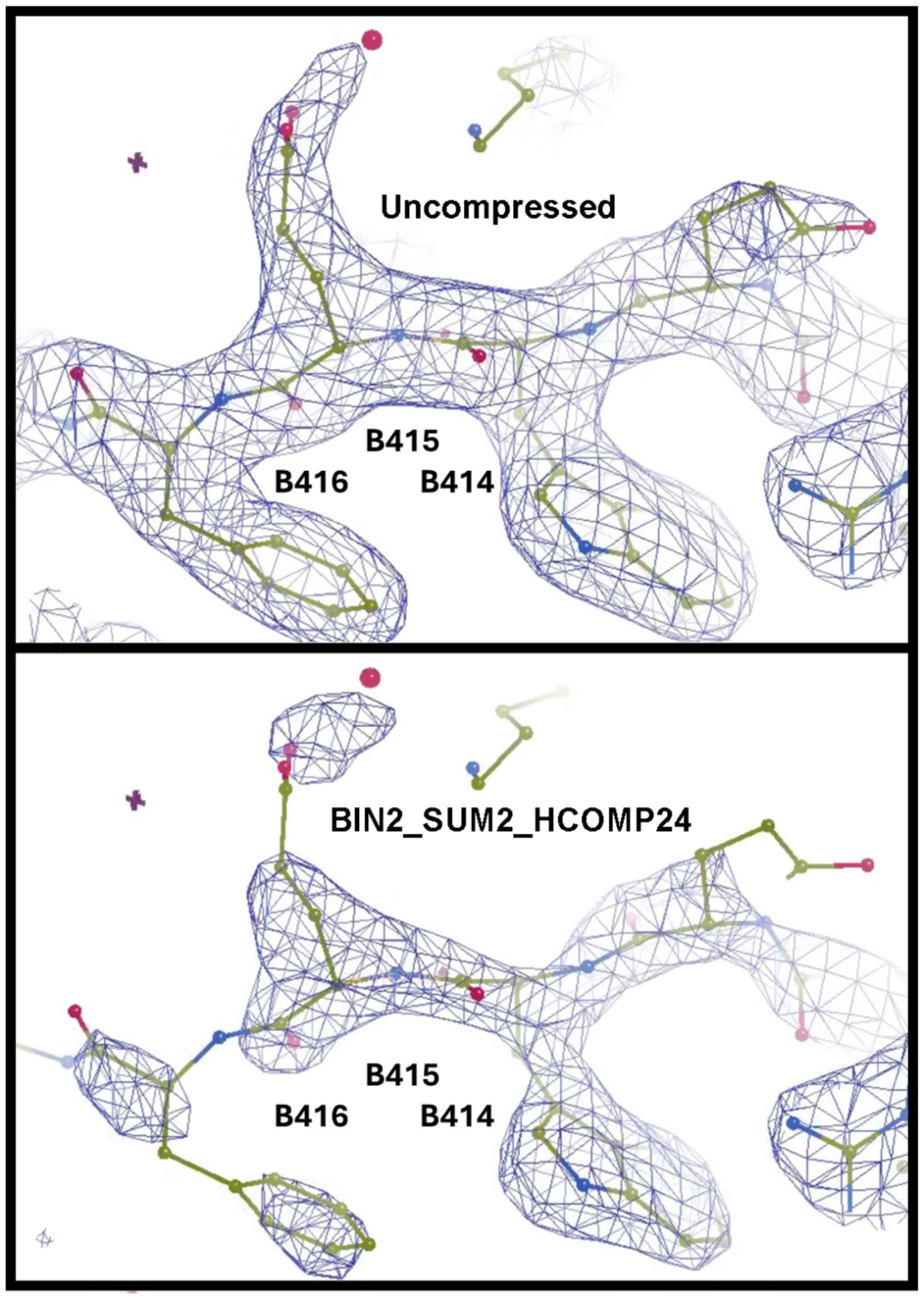
Electron density is shown for a region that contains some amino acids where the fit to the compressed data is good (right, B_414_), moderate (middle, B415), and poor (left, B416). Compare the electron density in the omit region obtained using the uncompressed data (top image) and those obtained using the BIN2_SUM2_HCOMP24 data (bottom image), which has a compression ratio of 627. Electron density is shown for a region that contains some amino acids where the fit to the compressed data is good (right, B_414_), moderate (middle, B_415_), and poor (left, B_416_). This region was selected specifically to demonstrate that (i) over-compression can initially manifest as difficult to detect locally poor fit between model and data, affecting only a few amino acids, and (ii) very aggressive compression ratios should be avoided in cases where good model-to-data fit is required for all amino acids in a structure.

**Figure 6.**
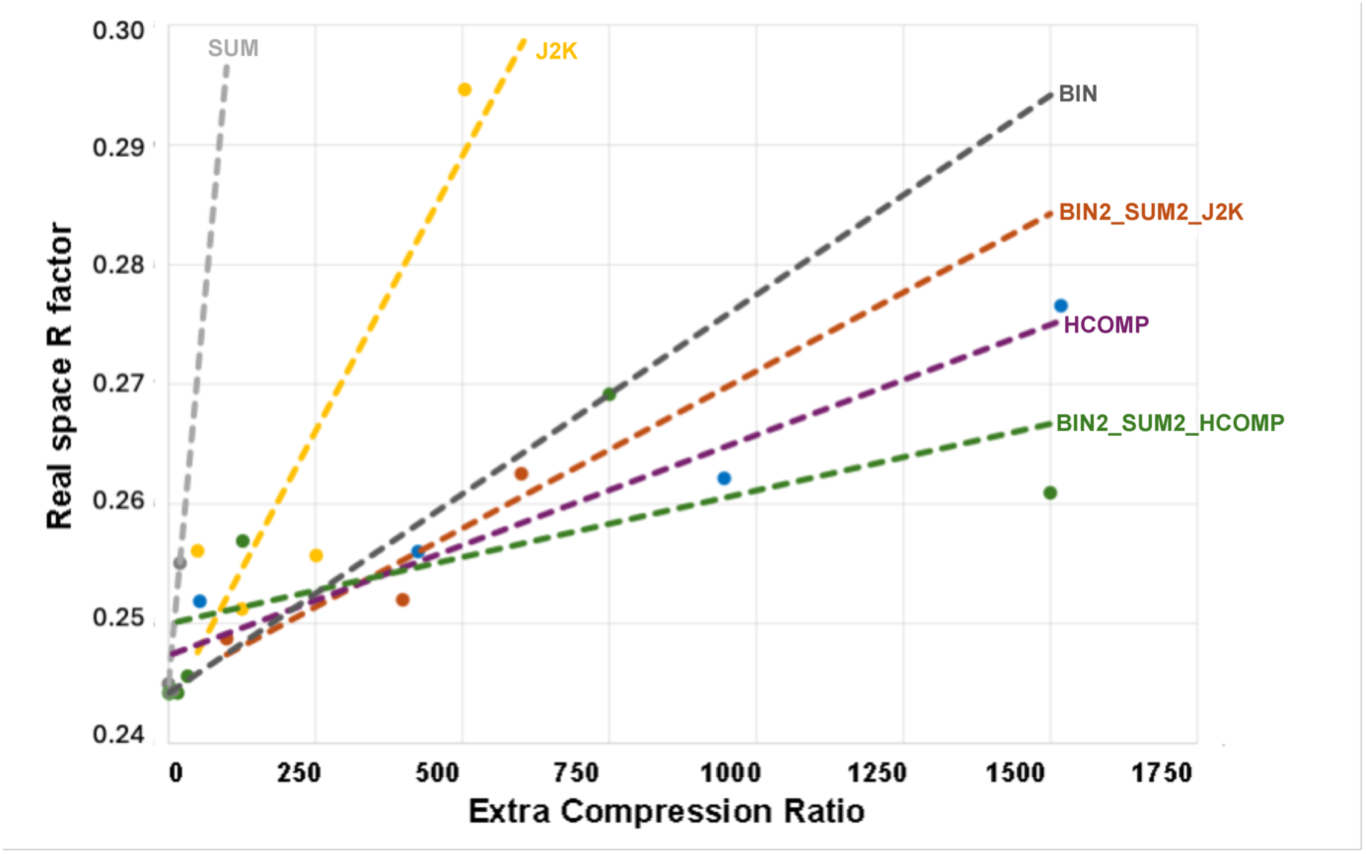
Benchmarking the effectiveness of compression strategies using real space R values for HIV reverse transcriptase.. Real space R values were computed (i) using *overlapmap* and (ii) using custom software (the quotient in the former case is the total volume electron density, while in the latter case it is the variance in the electron density). The results of these two strategies were almost identical (see Tables S6 and S7); the average of the two measurements is plotted (Y axis) as a function of the compression ratio (X axis). It is evident from the image that increasing compression ratios are associated with decreasing map quality (increased real space R value), and *the slope that governs this loss of quality is different for different compression strategies* (the lower the slope, the better the compression strategy). In the case of the data examined here the optimal strategy is to combine the effects of multiple compression strategies. Binning two pixels in all three measured reciprocal space dimensions (x, y, and phi) followed by using the compression algorithm that was developed for astronomy purposes (this is labelled BIN2_SUM2_HCOMP above).

**Figure 7.**
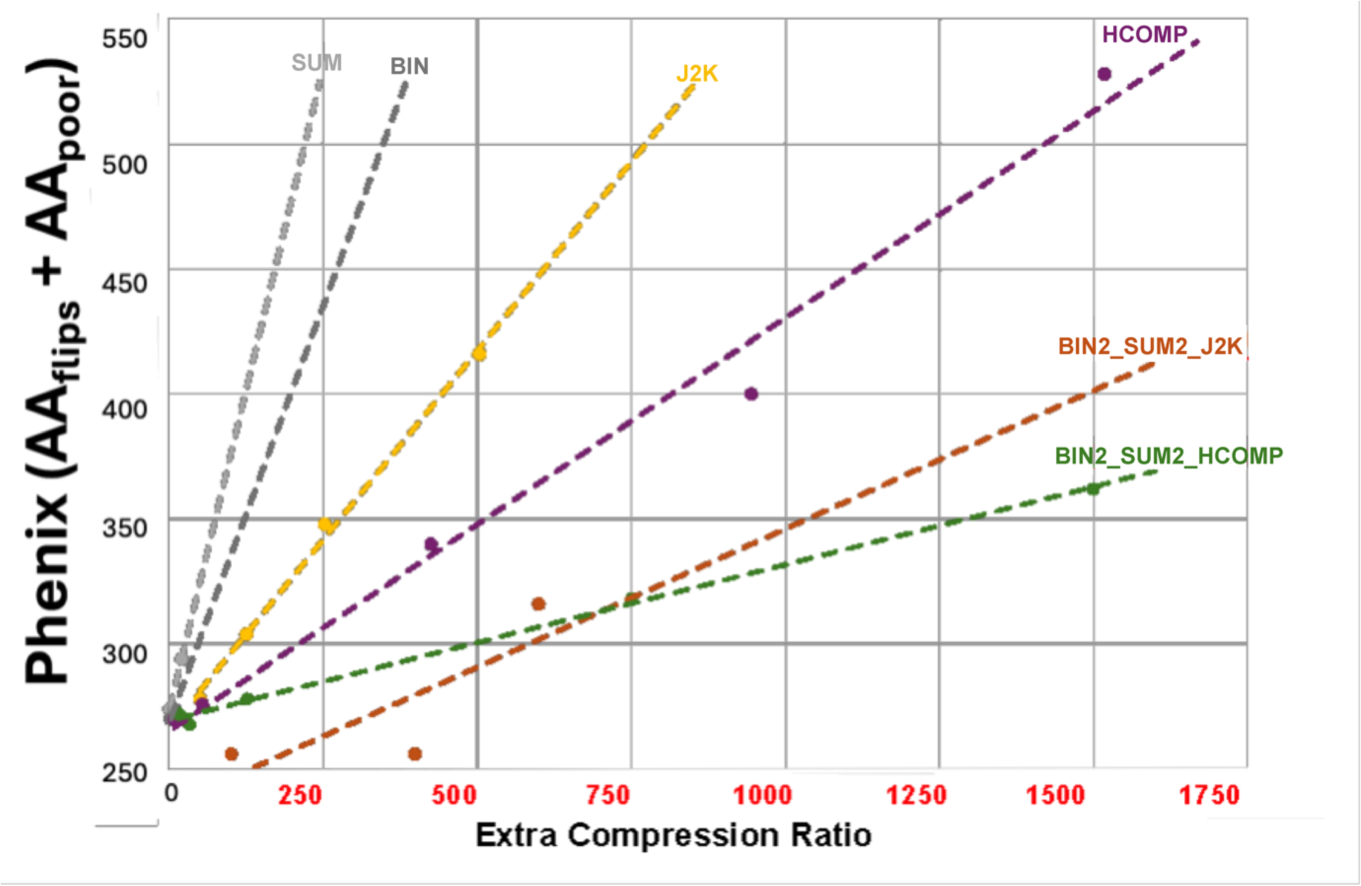
Benchmarking the effectiveness of compression strategies for HIV reverse transcriptase using binary metrics. The software package PHENIX was used to compare the electron density obtained from uncompressed data with the electron density obtained from data that were compressed using various strategies. The binary metrics (i) poor fit and (ii) conformational flip were added to indicate the total number of amino acids that were flagged as incorrect (hence the Y axis shows the number of amino acids that did not fit the observed electron density <u>plus</u> the number of amino acids that only fit with a different conformation). Just as in the case of Fig. 6, the slope of the decrease in quality varies between different compression strategies (and a smaller slope indicates a better compression strategy). Significantly, the best overall performance (labelled BIN2_SUM2_HCOMP) was the same when benchmarked using real space R values and when benchmarked using binary metrics.

An **HIV Reverse Transcriptase Workflow** is shown in the supplementary material. The workflow was used to benchmark the effect of data compression on HIV RT data and on the resulting model (including a large 188 amino acid omit region).

In order to compute real space R-values in a way that (i) permitted the comparisons of maps obtained using non-cognate atomic coordinates and (ii) permitted a custom definition of the real space R-value, such that the denominator was the variance of the reference map, rather than the total integrated electron density, we apply the algorithm as shown in supplemental Fig. S1.

In general experimental phasing projects do not require an initial pdb model. On the other hand ligand binding and MR projects do need an initial pdb model, and electron density maps are biased toward the model. For these experiments, one can afford higher compression ratios, compared to experimental phasing requiring more accurate reflection intensities.

Reducing model bias is a well-established challenge in MX structures that are solved using molecular replacement (Ramachandran *et al*., 1961) (Wang *et al*., 1985). While it is tempting to presume that the signal-to-noise advantage would allow higher compression ratios of molecular replacement experiments compared to experimental phasing experiments, we should be careful to ensure that model bias problems are not excessively increased at high compression ratios. To this end, we omitted a large domain from the model (188 residues between Thr^B^_240_ to GLN^B^_428_) before phasing each of the data sets that were compressed and then decompressed. We then were able to separately compute model correctness statistics (Table S8) and model to data fit statistics (Tables S9, S10, and S11 in the supplementary material) for the entire protein and for the omitted region. We used these benchmarking results to quantify the impact of lossy data on model bias and to establish reasonable limits for prudent compression of primary data. Our results indicated that the electron density map obtained after molecular replacement remains reasonably correct in most regions when data are compressed by a factor of 500, even in the case of marginal data (although some errors may appear in large, omitted regions; see Fig. 5).

Table 2 shows statistics for two very conservative extra compression runs for HIV reverse transcriptase of up to 400:1. Tables S6 and S7 in the supplementary material show more aggressive extra compression runs of as much as 1000:1 with satisfactory results.

**Table 2.** Sampling of successful HIV reverse transcriptase processing of compressed data, which worked for lossy compressions of more than 200:1. The values in parentheses are for the highest resolution shell.

| Compression | Extra Cmp Ratio | Rm all | Rpim all | Mean I/$\sigma$(I) | Compl | Tot Obs Unique |
| --- | --- | --- | --- | --- | --- | --- |
| uncompressed | 1 | 7.9 (144.8) | 4.1 (72.9) | 11.4 (1.0) | 99.9 (100.0) | 41196 (4602) |
| BIN2_SUM2_J2K50 | 200 | 11.7 (273.1) | 6.0 (137.3) | 8.1 (0.6) | 99.9 (100.0) | 41161 (4586) |
| BIN2_SUM2_J2K100 | 400 | 15.1 (319.4) | 7.8 (161.0) | 7.3 (0.5) | 99.9 (100.0) | 40963 (4557) |

## 4. Discussion

### 4.1. Hcompress vs. JPEG-2000, Impact of Reflection Profiles

The combined use of modest degrees of binning and summing strengthens the weak reflections, which helps to protect them from the impact of the massive compressions provided by Hcompress or JPEG-2000. See Figs. 8 -- 11 for the impact of JPEG-2000 and Hcompress by themselves and in combination with binning and summing by two when working with the HIV reverse transcriptase data.

**Figure 8.**
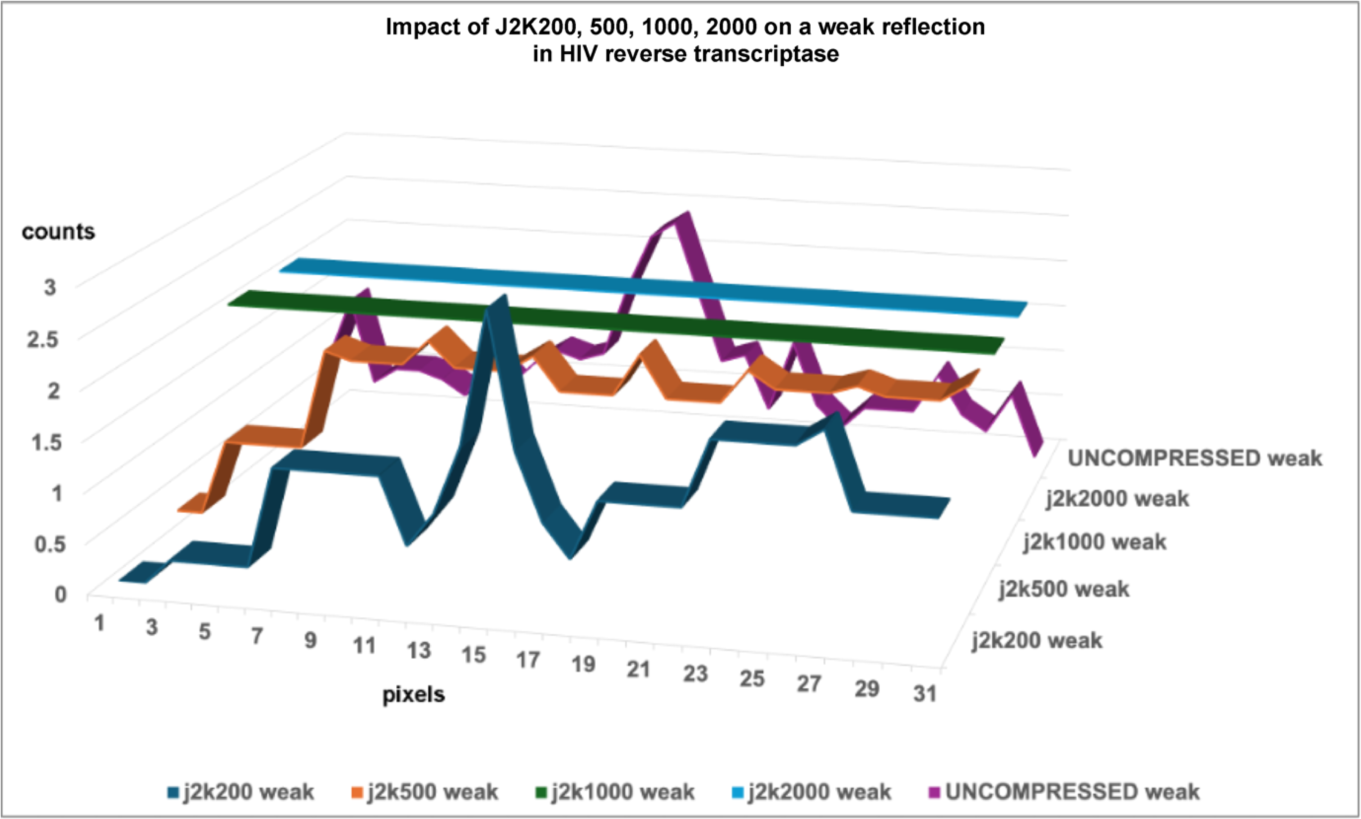
Impact of JPEG-2000 compressions at levels 200, 500, 1000 and 2000 on a typical weak reflection in HIV reverse transcriptase. Six rows through the reflection were averaged. The j2k200 compression preserves the peak but distorts the shoulders asymmetrically. The j2k500 compression severely distorts the profile, while j2k1000 and j2k2000 swallow the weak reflection completely.

Fig. 8 shows the impact of JPEG-2000 compressions on weak reflections in HIV reverse transcriptase. J2K200 keeps the weak peaks findable but distorts the shoulders. The higher JPEG-2000 compressions do serious damage, losing the peak entirely for J2K1000. Fig. 9 shows that Hcompress does better but also loses the peak entirely for high compressions. In Figs. 10 and 11 the weak peak has been maintained by modest binning and summing thereby allowing useful extra compression ratios well over 250:1 to be achieved in many cases and over 500:1 in some cases.

**Figure 9.**
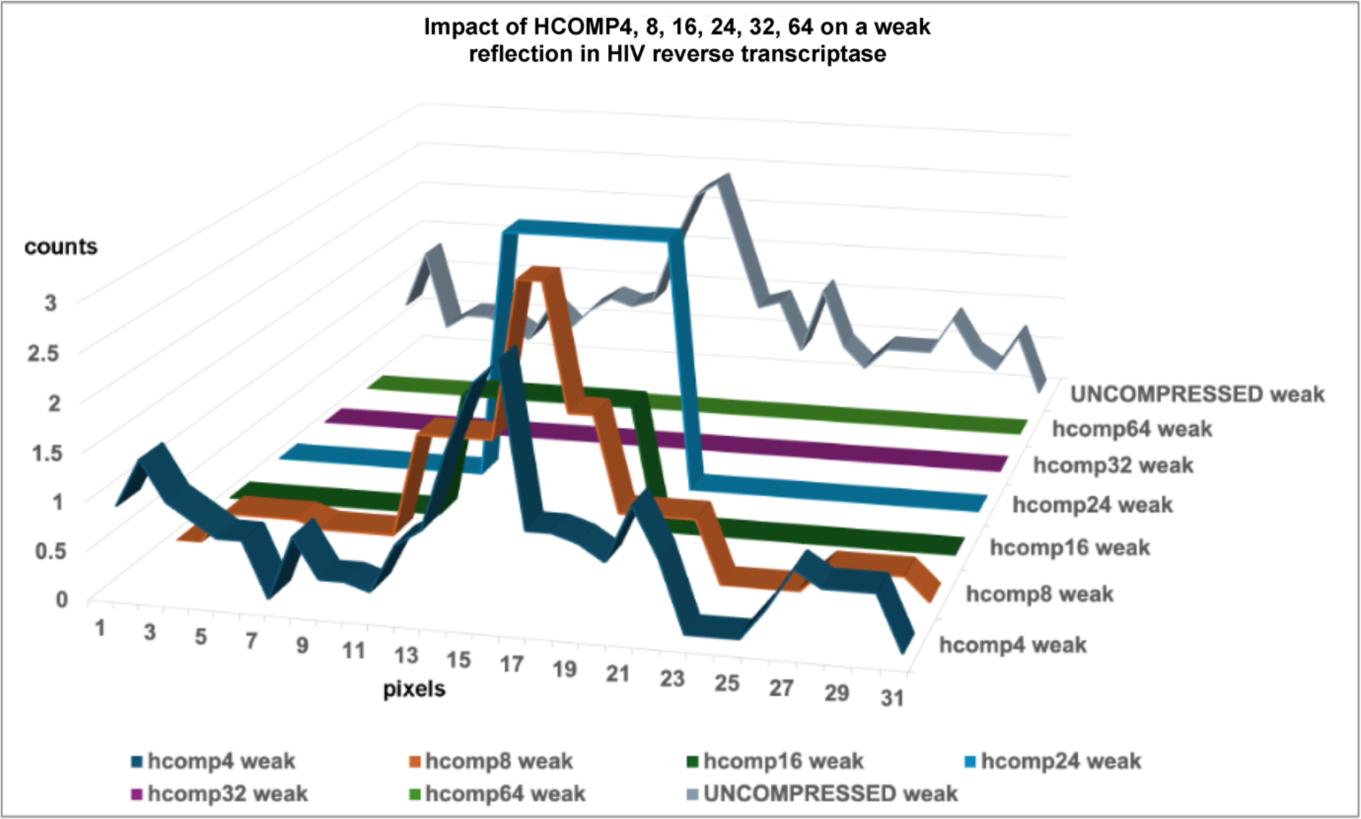
Impact of Hcompress compressions at scales 4, 8, 16, 24, 32 and 64 on a typical weak reflection in HIV reverse transcriptase. Six rows through the reflection were averaged. The hcomp 4, 8 and 16 compressions preserve the peak, but the hcomp16 peak has been distorted. The hcomp24 compression actually boosted this weak reflection and hcomp32 and 64 swallowed it.

**Figure 10.**
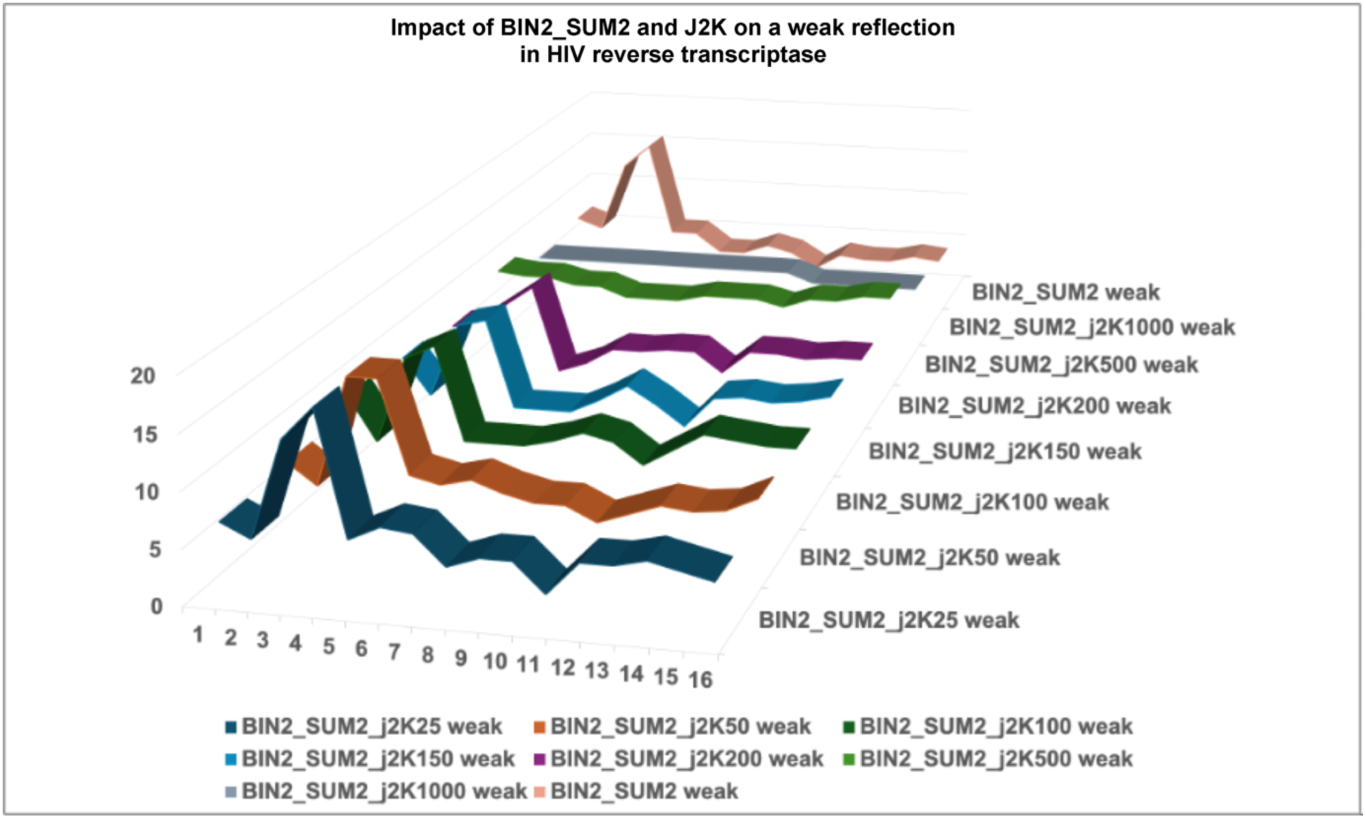
Impact of JPEG-2000 compressions at levels 25, 50, 100, 150, 200, 500, and 1000 on a typical weak reflection in HIV reverse transcriptase, first boosted and protected by binning and summing by factors of 2. Three rows through the boosted reflection were averaged. The BIN2_SUM2_J2K25, 50, 100, 150, 200 and 500 all preserve the basic shape of the weak reflection with some weakening that preserves the peak but distorts the shoulders asymmetrically. The j2k500 compression severely distorts the profile, while j2k1000 and j2k2000 swallow the weak reflection completely.

**Figure 11.**
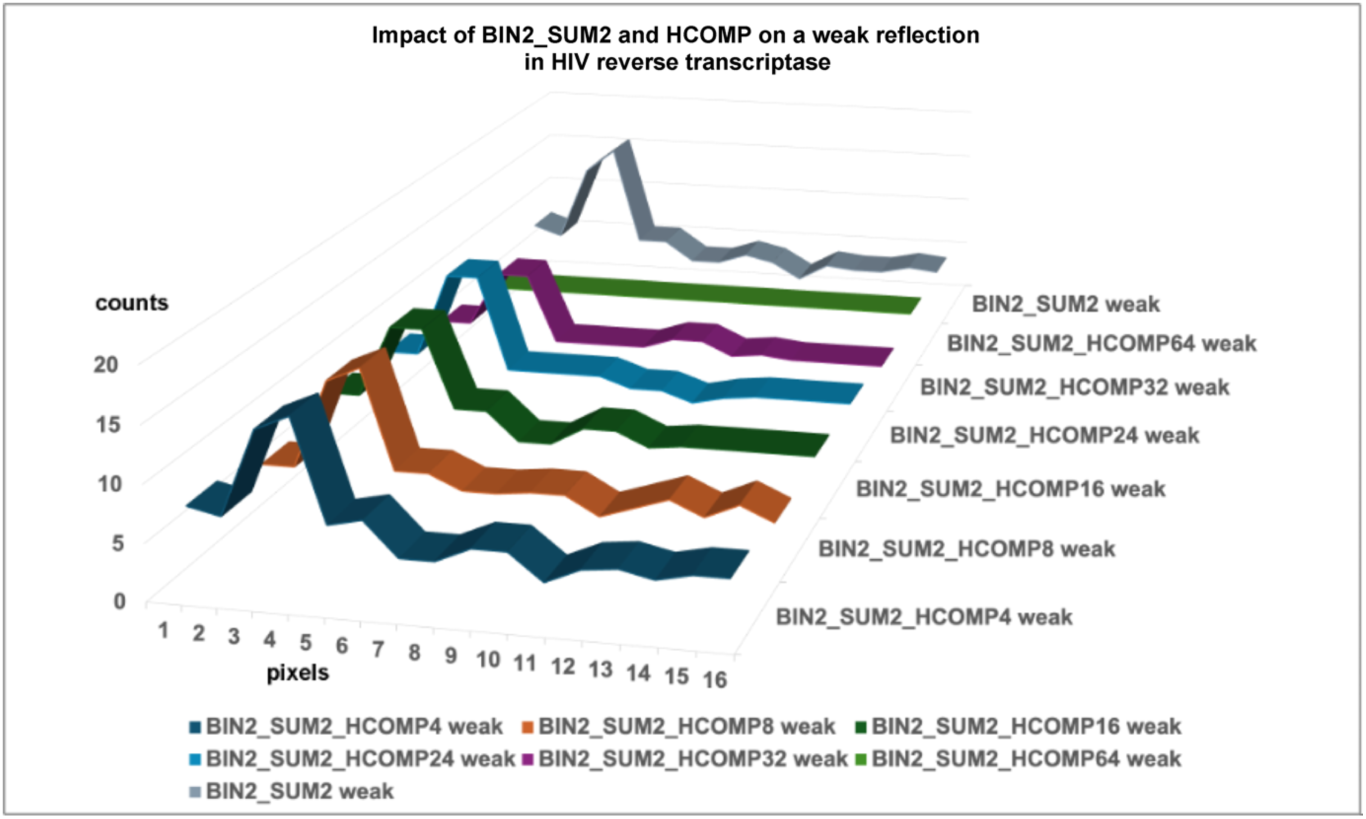
Impact of Hcompress compressions at scales 4, 8, 16, 24, 32, and 64 on a typical weak reflection in HIV reverse transcriptase first boosted and protected by binning and summing by factors of 2. Three rows through the boosted reflection were averaged. The BIN2_SUM2_HCOMP 4, 8, 16 and 24 all preserve the basic shape of the weak reflection with no weakening or distortion. The HCOMP32 compression weakens the reflection slightly. The HCOMP64 compression swallows the weak reflection completely.

The results we present reveal that with care to protect weak reflections with modest binning and summing, and with care to avoid over-compression, in many cases massive lossy compression can be applied successfully while retaining the information necessary to find and integrate Bragg reflections and to solve structures with good statistics.

### 4.2. Suggestions for Real Time Compression of MX Data

We expect that new detectors with substantially higher framing rates combined with synchrotron-source upgrades and expected new sample-delivery methods, will expectedly require the implementing of lossy compression. For many of the upcoming experiments, it will certainly be possible to engage lossy compression for real time processing depending on the type of experiment.

Using our initial results, one can extrapolate effects of binning (4x for 2x2) combined with either JPEG-2000 or Hcompress for real time lossy compression. We would suggest that one consider BIN2_HCOMP32 or BIN2_JPEG125 with compression ratios of 750 or 500 for medium intensities and resolution, as for the HIV reverse transcriptase data set, and 150 or 250 for strong intensities and high resolution, as for the lysozyme data set. These compressions offer minimum loss of resolvable structural information regardless of the experiment. For greater compression ratios, we suggest BIN2_HCOMP64 offering compression ratios of 250 and 1200 for the strong and medium intensities.

### 4.3. Suggestions for Long Term Data Archiving

For archiving of data, one can achieve significantly higher compression ratios by combining these three: pixel binning (4x for 2x2), image summing (2x, 4x, 5x) and subsequent image compression using either HCOMP64 or J2K125. One can achieve compression ratios of up to 500 to 1000 depending on the experiment. In other words, one can archive three years of a beamline operation (600 days) in the space originally required for ten to twenty hours of data throughput. Highlighted in this way, it puts the compression discussion into a broader perspective of archiving data “forever”.

## 5. Availability of Data and Software

The lossless compression data for the lysozyme experiment have been deposited in Zenodo^7^ and the scripts in GitHub^8^. PDB depositions have been made for samples of the uncompressed and compressed structures of lysozyme. The entry from the uncompressed data is 9B7F. The entry for the data compressed with BIN2_CBF2_HCOMP24 is 9B7E.

## 6. Over-compression

While a surprisingly high degree of lossy compression that impacts the background and not the Bragg peaks is close to harmless for many macromolecular crystallographic experiments, there are risks in overdoing lossy compression. If the compression reaches into the pixels under the Bragg peaks or disturbs the relationship between the local background level and the Bragg peaks, peaks may be lost or “accidentally inflated” in intensity. In the special case where diffuse scattering data is needed, requiring data from between the Bragg peaks, the best plan is both to keep some percentage of images (or possibly some percentage of complete data sets) faithfully compressed and to limit the level of lossy compression used on the remaining images to be within well-tested limits derived from processing similar structures.

Every lossy compression technique has limits beyond which image statistics are severely degraded. It is prudent to avoid “over-compression” and stay well within those limits. See the examples marked with yellow backgrounds in the supporting information.

## 7. Summary and Conclusions

We employed three existing and well-studied methods to accomplish dramatic compression where some information was lost, but much information was retained: binning of pixels, summing of frames, and JPEG-2000 from the movie industry or Hcompress from astronomy. For most MX images, one can almost always achieve at least 2:1 lossless compression, as provided by the detector control unit. The combination of 2x2 binning, summing by 2, and Hcompress was particularly effective in achieving useful extra lossy compressions of between 250:1 and 500:1 when the primary focus of the experiment is to process Bragg-refection data. As a result, one can reliably reduce the storage by a factor of better than 300:1. This will allow us to be sure to save enough data to be able to detect cases for which non-Bragg background data will need to be processed, so that very effective compression ratios will be achievable for the transition to much faster detectors. In this discussion we are not addressing throughput but are focusing on achievable compression ratios and the effect of the compressions used on the quality of the structures produced.

These results encourage us to study lossy compression further using crystallographic knowledge to subtract the whole background. In many cases we would compress only pixels directly associated with Bragg peaks. That will allow one to preserve important experimental data with overall compression ratios of between 500:1 and 10,000:1 in many high-data-rate macromolecular experiments.

## 8. Recapitulation Discussion

Lossy compression techniques can significantly reduce data size by focusing on retaining essential properties while sacrificing less critical ones. Key questions to address are: (i) which data properties should be preserved, (ii) which compression methods best maintain these properties, and (iii) how to collect data to achieve high compression ratios. Future work should explore these questions, such as developing better diffuse scattering techniques, matching compression strategies to specific data types, and optimizing data collection methods. This discussion aims to ignite a dialogue on lossy compression for MX data, recognizing that it may affect different community members unevenly and valuing diverse perspectives.

## Acknowledgements

We gratefully acknowledge thoughtful comments and suggestions by Robert M. Sweet and Frances C. Bernstein.

The entire community owes a debt to James Holton for his creative, useful, and constructive comments and suggestions on the subject of lossy compression for macromolecular crystallography since 2011. He helped us all to break free of the past and look dispassionately at what modern realities require.

We thank Dr. Karen Anderson for generously allowing us to use her HIV reverse transcriptase data to benchmark our compression algorithms, and for her comments regarding the ways that small changes in data quality could affect the interpretability of data in the vicinity of key areas of the protein that are biologically significant.

This work utilized the AMX beamline from the Center for BioMolecular Structure (CBMS) which is primarily supported by the National Institutes of Health, National Institute of General Medical Sciences (NIGMS) through a Center Core P30 Grant (P30GM133893), and by the DOE Office of Biological and Environmental Research (KP1607011). As part of NSLS-II, a national user facility at Brookhaven National Laboratory, work performed at the CBMS is supported in part by the U.S. Department of Energy, Office of Science, Office of Basic Energy Sciences Program under contract number DE-SC0012704.

## Supporting information

### Worked Example of Estimated Costs at AMX

*With current beamline performance and upgraded detectors and either assuming very efficient users, or, better, users relying mostly on automated data collection and screening, at AMX, one can achieve 700 samples per day. With each sample generating about 2-5 GB of raw diffraction data per data set and assuming that 2-3 data sets are collected per sample, per standard and per vector, this corresponds to 10 GB per sample, and 7 TB / day. Note that for each sample, using autoProc and fast dp, directories are created in data processing with 0.5 TB of files. Each sample also undergoes rastering with on average 0.5 TB per sample. Subsequent data analysis may generate 0.5 TB of data too*.

*Only raw diffraction, metadata, data reduction final log and reduced files as well as structure files should be kept along with input and instruction files. Rastering data and intermediate files can be deleted*.

*This corresponds to up to 11 GB per sample, or 7.7 TB per day and 1.2 PB / year per highly productive and efficient beamline. This is calculated based on 5000 operating hours and 75 % of total time used for user operation*.

*Improvements in throughput will likely result in double or triple that amount in the next 5 years*.

*Slow access Amazon Web Services data (Amazon glacier) costs 0.001 cts / GB / month : i.e. per beamline US$1200 per month or US$14,400 per year. This excludes added ingress and egress costs to retrieve and download or upload data*.

*Assuming that a compression ratio of 1000 can be achieved, one would need to archive 1.2 TB / year. A dual set of RAID1 arrays with a capacity of 4 TB using SSD drives costs ∼ US$1500, not counting labor costs*.

*The more important cost saving is related to facility wide storage: ∼ US$200,000/ year for a 4 PB file system maintained for 5 years*.

*Big question is the computing cost to run these: beam down time: more or less no cost*.

## Supplementary Tables

Additional compression and structure determination runs were performed for both lysozyme, as shown in Tables S1, S2, S3, S4, and S5, and HIV reverse transcriptase, as shown in Tables S6 and S7. The Table heading arp_100_res_Cycle is the round at which arp built 100 residues or more for lysozyme. The areas of the Tables that are bolded indicate good runs in structure determination. The areas with light blue background required use of heavy atoms from a different compression method or otherwise showed data degradation. The areas with a yellow background indicate runs with serious issues in structure determination.

**Table S1.** Lysozyme compression runs. The successful runs are bolded.

| Extra Compression Ratio | Compression | SHELXD Sol | SHELXE RES | SHELXE FOM/ CONN | DM FOM | ARP res / correct | R | arp_100_res_Cycle; |
| --- | --- | --- | --- | --- | --- | --- | --- | --- |
| 1 | **CBF01 (none)** | 17 | **119** | **0.65/0.77** | **0.83** | **127/1.00** | **22.4** | **2 (110)** |
| 25 | **J2K50** | 8 | **111** | **0.65/0.76** | **0.83** | **127/1.00** | **25.4** | **2 (103)** |
| 50 | **J2K100** | 4 | **113** | **0.61/0.76** | **0.82** | **127/1.00** | **23** | **3 (102)** |
| 100 | **J2K200** | 4 | **104** | **0.65/0.76** | **0.82** | **127/1.00** | **23.1** | **3 (105)** |
| 251 | J2K500<br>J2K500 - HA_solutio<br>n-J2K200 | 0<br>NA | 0<br>110 | 0<br>0.64/<br>0.76 | 0<br>0.81 | 0<br>127/<br>1.00 | 0<br>25.3 | 5 (101) |
| 498 | J2K1000<br>J2K1000 - HA_solutio<br>n-J2K200 | 0<br>NA | 0<br>59 | 0<br>0.56/<br>0.71 | 0<br>0.73 | 0<br>127/<br>1.00 | 0<br>28.9 | 15 (105) |
|  | J2K1000:<br>10000trials |  |  |  |  |  |  |  |
| 748 | J2K1500 | NA | NA | NA | NA | NA | NA |  |
|  | no HA solution; tried 1 2 and 5k; using HA peaks from jpg200; no success either |  |  |  |  |  |  |  |
| 995 | J2K2000 | NA | NA | NA | NA | NA | NA |  |
|  | re-ran:minvalpix=2; forced cell; poor stats |  |  |  |  |  |  |  |
| 8 | **BIN2_**<br>**SUM2** | 6 | **109** | **0.63/0.76** | **0.83** | **127/1.00** | **23.3** | **3 (102)** |
| 100 | **BIN2_**<br>**SUM2_**<br>**J2K25** | 16 | **109** | **0.65/0.77** | **0.83** | **127/1.00** | **23.4** | **2 (111)** |
| 200 | **BIN2_**<br>**SUM2_**<br>**J2K50** | 5 | **116** | **0.64/0.76** | **0.83** | **127/1.00** | **23.4** | **2 (106)** |
| 399 | **BIN2_**<br>**SUM02_**<br>**J2K100** | 16 | **106** | **0.64/0.76** | **0.82** | **127/1.00** | **22.9** | **2 (100)** |
| 499 | **BIN2_**<br>**SUM02_**<br>**J2K125** | 8 | **105** | **0.64/0.76** | **0.82** | **125/1.00** | **24.4** | **2 (101)** |

**Table S2.** Lysozyme compression runs, continued. The successful runs are bolded.

| Extra Compression Ratio | Compression | SHELXD Sol | SHELXE RES | SHELXE FOM/CONN | DM FOM | ARP res / correct | R | arp_100_res_Cycle; |
| --- | --- | --- | --- | --- | --- | --- | --- | --- |
| 529 | **BIN2_SUM02_J2K130** | **1** | **110** | **0.63/0.76** | **0.82** | **127/1.00** | **23.6** | **2 (107)** |
| 559 | BIN2_SUM02_J2K140 | NA | NA | 0.58/NA | 0.77 | NA | 22.98 | NA |
| 600 | BIN2_SUM02_J2K150 | same destroys all the data |  |  |  |  |  |  |
| 797 | BIN2_SUM02_J2K200 | 18 (CC=10) | no phases |  |  | 10 res/0.66 con |  |  |
|  | with good peaks: no phases<br>jpeg 200 "destroys" data in ways that phasing does not work with good heavy atoms |  |  |  |  |  |  |  |
| 18 | **HCOMP4** | **10** | **105** | **0.64/0.76** | **0.82** | **127/1.00** | **24.5** | **2 (103)** |
| 96 | **HCOMP8** | **30** | **116** | **0.62/0.76** | **0.8** | **125/1.00** | **25.8** | **2 (104)** |
| 320 | **HCOMP16** | **0.8 (1/1200)** | **102** | **0.65/0.76** | **0.81** | **127/1.00** | **23** | **2 (104)** |
| 335 | **HCOMP20** | **0.8 (1/1200)** | **102** | **0.66/0.76** | **0.81** | **127/1.00** | **23.9** | **2 (102)** |
| 357 | **HCOMP24** |  | **106** | **0.62/0.77** | **0.82** | **127/0.998** | **24.2** | **4 (104)** |
|  | 0/10000 (usedhcomp08_HA)) |  |  |  |  |  |  |  |
| 480 | **HCOMP32** | **5/5000 (>2500)** | **117** | **0.67/0.76** | **0.8** | **127/1.00** | **22.4** | **2(115)** |
| 672 | **HCOMP64** | **0/5000** | **32** | **0.56/0.74** | **0.72** | **5** | **20.9** | **NA** |
| 17 | **BIN2_SUM02_HCOMP4** | **2** | **106** | **0.65/0.76** | **0.83** | **127/1.00** | **23.5** | **2 (108)** |
| 36 | **BIN2_SUM02_HCOMP08** | **11** | **109** | **0.60/0.76** | **0.82** | **127/1.00** | **23.6** | **5 (101)** |
| 112 | **BIN2_SUM02_HCOMP16** | **15** | **103** | **0.63/0.76** | **0.82** | **127/1.00** | **23.1** | **4 (102)** |
| 211 | **BIN2_SUM02_HCOMP24** | **14** | **99** | **0.61/0.76** | **0.82** | **128/0.999** | **26.7** | **6 (122)** |

**Table S3.** Lysozyme compression runs, continued. The successful runs are bolded.

| Extra<br>Compression<br>Ratio | Compression | SHELX<br>D Sol | SHELXE<br>RES | SHELXE<br>FOM/<br>CONN | DM<br>FOM | ARP res<br>/ correct | R | arp_100_<br>res_Cycle; |
| --- | --- | --- | --- | --- | --- | --- | --- | --- |
| 322 | **BIN2_SUM02_**<br>**HCOMP32** | **10** | **112** | **0.65/**<br>**0.76** | **0.81** | **127/**<br>**1.00** | **23.7** | **3 (102)** |
| 457 | **BIN2_SUM02_**<br>**HCOMP64** | **1** | **105** | **0.63/**<br>**0.76** | **0.79** | **127/**<br>**1.00** | **24** | **2 (100)** |
| 10 | **BIN2_SUM05** | **7** | **109** | **0.66/**<br>**0.77** | **0.84** | **127/**<br>**1.00** | **22.4** | **1 (105)** |
| 62 | **BIN2_SUM05_**<br>**HCOMP4** | **0.15** | **112** | **0.65/**<br>**0.77** | **0.83** | **127/**<br>**0.999** | **23.5** | **2 (102)** |
| 220 | **BIN2_SUM05_**<br>**HCOMP16** | **12** | **95** | **0.63/0.7**<br>**6** | **0.82** | **127/**<br>**0.999** | **23.7** | **3 (106)** |
| 1185 | **BIN2_SUM05_**<br>**HCOMP64** | **5** | **111** | **0.62/0.7**<br>**5** | **0.8** | **126/**<br>**0.999** | **24.2** | **2 (110)** |
| 20 | **BIN2_SUM10**<br>4 / 2000 (1185) | **0.84** | **108** | **0.64/0.7**<br>**7** | **0.83** | **127 /**<br>**1.00** | **22.8** | **3 (108)** |
| 100 | **BIN2_SUM10_**<br>**HCOMP4**<br>10 / 10 000 (2329) | **0.43** | **106** | **0.64/0.7**<br>**6** | **0.83** | **127 /**<br>**1.00** | **22.9** | **2 (101)** |
| 256 | **BIN2_SUM10_**<br>**HCOMP16** | **1** | **115** | **0.63/0.7**<br>**6** | **0.83** | **127 /**<br>**1.00** | **23.8** | **2 (112)** |
| 1336 | **BIN2_SUM10_**<br>**HCOMP64** | **2** | **94** | **0.64/0.7**<br>**5** | **0.79** | **127 /**<br>**1.00** | **23.9** | **4 (109)** |

Lysozyme data reduction was done in an overall range of 19.72 Å to 1.64 Å with the highest resolution shell being 1.67 Å to 1.64 Å. All data were processed using the same xds.inp that was adjusted for binning and by 2s (pix size, beam center, osc range and masked areas).

Tables S4 and S5 shows the results of the data reduction runs for all lysozyme data sets. Boldface indicates successful runs. A light blue background indicates runs with issues. A light yellow background indicates runs that failed. The Tables highlight the most commonly used parameters (Rmerge, Rpim, completeness, mean I/σ(I), unique reflections) as well as a subsequent set of parameters to illustrate potential damage to data statistics prior to S_SAD phasing (anomalous correlation and mid-slope anomalous probability). For simplification, other parameters usually displayed are not included in this Table. 3600 frames were collected at 7.5 keV to improve Sulfur dF” to maximize likelihood of successful phasing using anomalous signal from the 10 S atoms present in the lysozyme. Note that the protein was crystallized in a NaCl solution resulting in observation of 7 additional peaks that are either Na or Cl ions. The numbers in parentheses are from the highest resolution shell.

**Table S4.** Lysozyme data reduction runs. The successful runs are bolded. Tables S4 and S5 show the data reduction statistics for all lysozyme data sets.

| Compression | Rm all | Rpim all | Mean I/$\sigma$(I) | Completeness | Delano correl $\frac{1}{2}$ | Mid-slope anom prob | Tot Obs Unique |
| --- | --- | --- | --- | --- | --- | --- | --- |
| **native** | **4.1 (7.8)** | **0.9 (6.0)** | **60.9 (8.5)** | **88.9 (19.9)** | **0.427 (0.088)** | **1.842** | **13089 (140)** |
| J2K50 | 4.1 (9.8) | 1.0 (8.6) | 54.5 (7.6) | 87.6 (20.3) | 0.437 (0.126) | 1.7 | 12901 (154) |
| J2K100 | 4.6 (11.7) | 1.0 (8.8) | 50.9 (6.8) | 87.6 (19.7) | 0.405 (-0.315) | 1.593 | 12911 (139) |
| J2K200 | 4.8 (16.2) | 1.1 (12.4) | 47.1 (4.5) | 87.3 (19.7) | 0.355 (0.178) | 1.488 | 12859 (139) |
| J2K500 | 5.8 (33.4) | 1.3 (25.3) | 39.2 (2.0) | 87.2 (19.7) | 0.276 (0.370) | 1.297 | 12844 (139) |
| J2K1000 | 9.9 (47.4) | 2.2 (38.0) | 20.3 (2.2) | 84.5 (17.1) | 0.106 (-0.377) | 1.103 | 12475 (121) |
| J2K1500 | 24.2 (90.7) | 5.5 (73.9) | 8.3 (0.7) | 87.2 (17.8) | 0.043 (0.000) | 0.861 | 12707 (128) |
| forcing cell parameters (otherwise failed to converge) |  |  |  |  |  |  |  |
| J2K2000 | 125.3 (NA) | 71.5 (NA) | 0.6 (0.7) | 45.8 (2.4) | -0.147 (NA) | 0.73 | 6453 (16) |
| forcing cell parameters (otherwise failed to converge) |  |  |  |  |  |  |  |
| BIN2_SUM02 | 4.6 (7.8) | 1.0 (6.1) | 56.5 (8.5) | 88.7 (18.3) | 0.378 (0.214) | 1.758 | 13053 (129) |
| BIN2_SUM02_J2K25 | 4.5 (11.6) | 1.0 (9.2) | 55.4 (6.4) | 88.7 (18.3) | 0.376 (0.073) | 1.751 | 13059 (129) |
| BIN2_SUM02_J2K50 | 4.7 (17.3) | 1.0 (13.7) | 52.9 (4.8) | 88.7 (18.2) | 0.357 (-0.450) | 1.644 | 13058 (128) |
| BIN2_SUM02_J2K100 | 4.8 (23.4) | 1.0 (17.8) | 51.2 (3.9) | 88.7 (18.3) | 0.326 (-0.146) | 1.595 | 13062 (129) |
| BIN2_SUM02_J2K125 | 4.8 (28.6) | 1.0 (21.6) | 50.7 (2.9) | 88.7 (18.3) | 0.341 (0.179) | 1.553 | 13069 (129) |
| BIN2_SUM02_J2K130 | 4.8 (27.8) | 1.0 (21.3) | 50.4 (2.9) | 88.8 (18.3) | 0.343 (0.285) | 1.547 | 13070 (129) |
| BIN2_SUM02_J2K140 | 169.0 (208.4) | 36.3 (175.5) | 8.9 (0.6) | 88.1 (16.0) | 0.007 (-0.478) | 0.817 | 13065 (18) |
| BIN2_SUM02_J2K150 | 480.3 (-223) | 107.3 (-219.6) | 2.2 (-0.3) | 85.7 (4.0) | -0.092 (NA) | 0.716 | 12262 (32) |
| BIN2_<br>SUM02_<br>J2K200 | 994.7 (-<br>228.1) | 208.8 (-<br>228.1) | 1.9 (-0.1) | 83.1 (5.6) | -0.042<br>(NA) | 0.765 | 14201<br>(44) |
| HCOMP4 | 4.6<br>(9.2) | 1.0<br>(6.9) | 51.7<br>(7.6) | 88.8<br>(19.2) | 0.365 (-<br>0.066) | 1.696 | 13081<br>(135) |
| HCOMP8 | 5.1<br>(15.4) | 1.1<br>(11.9) | 44.6<br>(5.6) | 87.8<br>(17.0) | 0.369<br>(0.770) | 1.5 | 12911<br>(119) |
| HCOMP16 | 5.5<br>(16.3) | 1.2<br>(13.6) | 42.4<br>(4.8) | 84.8<br>(11.6) | 0.364<br>(0.000) | 1.477 | 12448<br>(81) |
| HCOMP20 | 5.7<br>(15.9) | 1.3<br>(13.0) | 41.2<br>(4.7) | 84.3<br>(11.1) | 0.330<br>(0.000) | 1.437 | 12366<br>(78) |

**Table S5.**
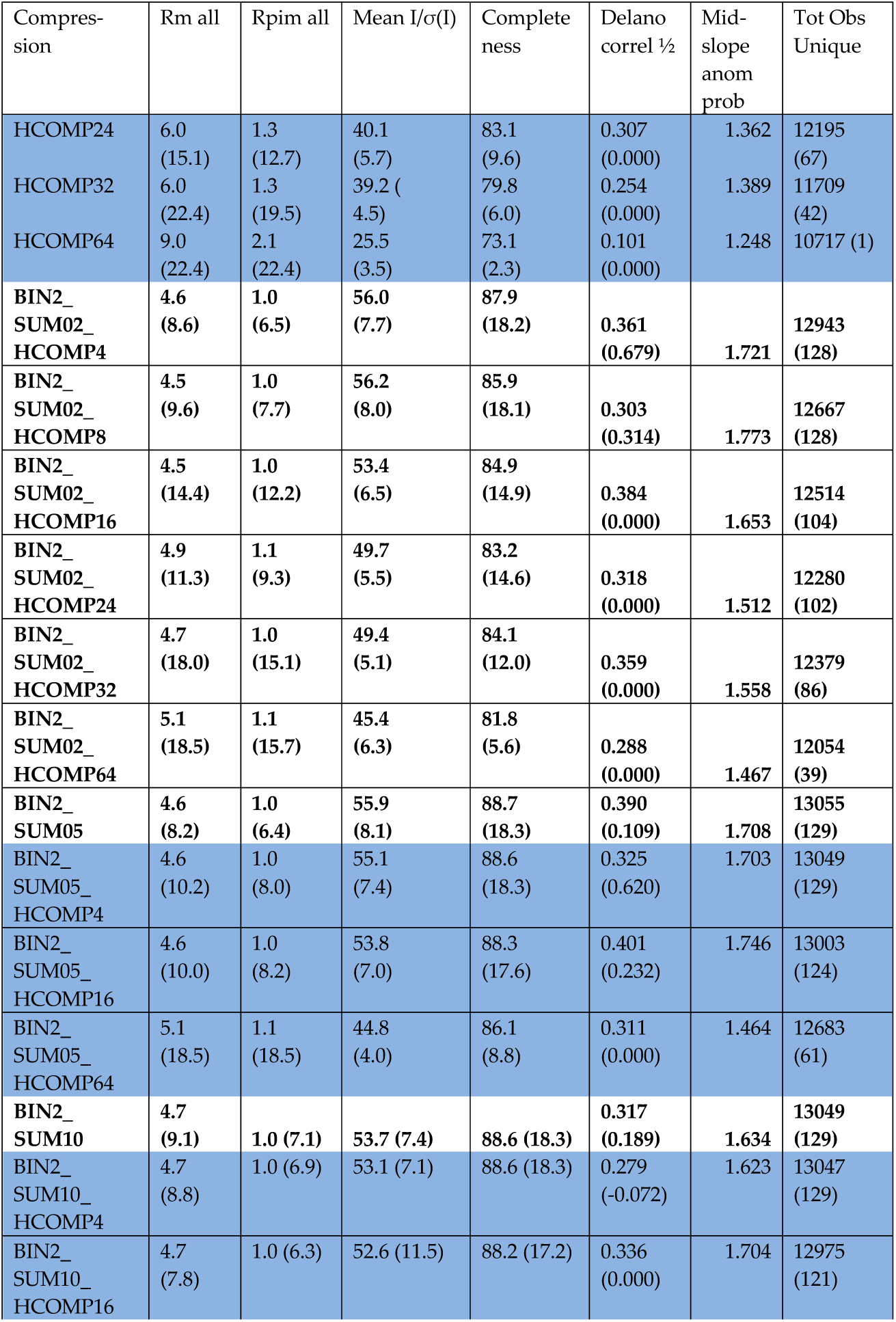
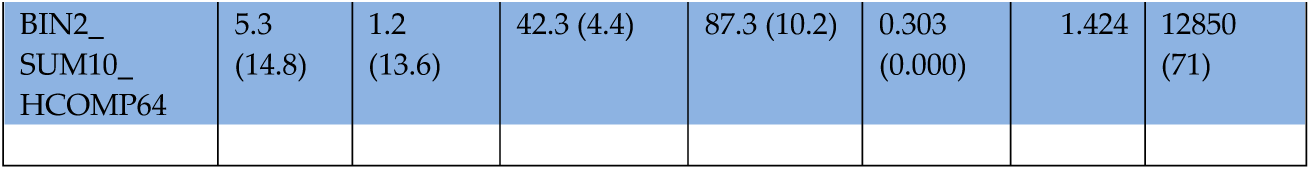
Lysozyme data reduction runs, continued. The successful runs are bolded. The numbers in parentheses are from the highest resolution shell.

### HIV Reverse Transcriptase Workflow

We developed a workflow to benchmark the effect of data compression on HIV RT data and on the resulting model (including a large 188 amino acid omit region). The following workflow was used:

1. HIV reverse transcriptase data were compressed and decompressed.
2. HIV reverse transcriptase were reduced using XDS.
3. An initial structure was obtained by the DIMPLE protocol, using a starting model with 188 residues omitted (from Thr^B^_240_ to GLN^B^_428_). The native data set was treated similarly but with no omit region.
4. The resolution of all data was reduced to 2.55 Å. The native data extended to 2.42Å.
5. Structures were refined using REFMAC and PHENIX. A single automated protocol was used for all data refinement.
6. Structure quality statistics were harvested from the atomic coordinate files (Table S8). Note that the large omit region resulted in increased R values.
7. The atomic coordinates resulting from the refinement against the native data (reduced to 2.55 Å) were copied to every directory containing compressed data. Note that this coordinate file did not have an omitted region.
8. The native structure was compared to each compressed data set to generate a real space R value using *overlapmap* (Table S9).
9. The native structure was compared to each compressed data set to generate a real space R value using custom software (Table S10, numbers on the right side).
10. The electron density map obtained using the native structure was then compared to the map obtained using each compressed data set to generate a real space R value using custom software (Table S10, numbers on the left side).
11. The native structure was compared to each compressed data set to generate model to data fit metrics using PHENIX (Table S11).
12. For steps 8 to 11, averages of the quality metrics were separately obtained for Chain A, and for the included and omitted regions of Chain B.

The data reduction for HIV reverse transcriptase is shown in Tables S6 and S7.

**Table S6.** HIV reverse transcriptase data reduction runs. The successful runs are bolded. The numbers in parentheses are from the highest resolution shell.

| Compression | Rm all | Rpim all | Mean<br>I/$\sigma$(I) | Completeness | Delano<br>correl $\frac{1}{2}$ |
| --- | --- | --- | --- | --- | --- |
| **Native** | **7.9**<br>(144.8) | **4.1**<br>(72.9) | **11.4 (1.0)** | **99.9**<br>(100.0) | **41196**<br>(4602) |
| **SUM02** | **8.4**<br>(161.9) | **4.3**<br>(81.3) | **11.1 (0.9)** | **99.9**<br>(100.0) | **41197**<br>(4601) |
| **SUM05** | **9.4**<br>(209.4) | **4.9**<br>(105.1) | **10.5 (0.7)** | **99.9**<br>(100.0) | **41217**<br>(4610) |
| **SUM10** | **11.1**<br>(269.6) | **5.8**<br>(136.2) | **9.3 (0.6)** | **99.8**<br>(100.0) | **41202**<br>(4613) |
| **SUM20** | **15.3**<br>(370.2) | **8.3**<br>(190.8) | **7.0 (0.4)** | **99.6**<br>(100.0) | **41106**<br>(4607) |
| **SUM40** | **21.3**<br>(578.4) | **12.7**<br>(312.9) | **5.2 (0.3)** | **98.4 (99.9)** | **40527**<br>(4600) |
| **BIN2** | **8.7**<br>(135.1) | **4.5**<br>(67.6) | **11.2 (1.1)** | **99.8**<br>(100.0) | **41184**<br>(4605) |
| **BIN4** | **12.2**<br>(155.1) | **6.3**<br>(77.1) | **9.4 (1.0)** | **99.7**<br>(100.0) | **41070**<br>(4590) |
| **BIN6** | **-6038**<br>(-745) | **-3105**<br>(-380) | **0**<br>(-0.2) | **91.1 (72.7)** | **37242**<br>(3313) |
| **J2K10** | **8.0**<br>(147.2) | **4.1**<br>(74.2) | **11.4 (1.0)** | **99.9**<br>(100.0) | **41197**<br>(4601) |
| **J2K20** | **9.1**<br>(198.1) | **4.7**<br>(99.9) | **9.7 (0.8)** | **99.9**<br>(100.0) | **41212**<br>(4600) |
| **J2K50** | **10.8**<br>(232.4) | **5.6**<br>(116.8) | **8.7 (0.7)** | **99.9**<br>(100.0) | **41210**<br>(4602) |
| **J2K75** | **12.3**<br>(265.5) | **6.3**<br>(133.5) | **8.8 (0.6)** | **99.9**<br>(100.0) | **41189**<br>(4597) |
| **J2K100** | **13.6**<br>(297.2) | **7.0**<br>(149.0) | **8.1**<br>(0.5) | **99.9**<br>(100.0) | **41204**<br>(4603) |
| **J2K150** | **15.1**<br>(341.0) | **7.8**<br>(170.5) | **7.6**<br>(0.4) | **99.9**<br>(100.0) | **41173**<br>(4594) |
| J2K200 | 23.8<br>(169.8) | 12.3<br>(87.8) | 6.0<br>(1.2) | 99.9<br>(100.0) | 41135<br>(4598) |
| J2K500 | 27.7<br>(399.5) | 14.3<br>(218.1) | 4.5<br>(0.4) | 99.6 (97.9) | 40996<br>(4497) |
| J2K1000 | 28.2<br>(-4756) | 14.7 (-<br>2416) | 3.6 (0.0) | 96.1 (85.5) | 39339<br>(3898) |
| J2K2000 | 81.9<br>(273.8) | 43.4<br>(162.8) | 1.6 (0.4) | 95.0 (94.7) | 39589<br>(4428) |

**Table S7.** HIV reverse transcriptase data reduction runs, continued. The successful runs are bolded. The numbers in parentheses are from the highest resolution shell.

| Compression | Rm all | Rpim all | Mean I/$\sigma$(I) | Completeness | Delano correl $\frac{1}{2}$ |
| --- | --- | --- | --- | --- | --- |
| **BIN2_SUM02** | **9.2**<br>(151.1) | **4.7**<br>(75.7) | **11.0**<br>(1.0) | **99.8**<br>(100.0) | **41164**<br>(4597) |
| **BIN2_SUM02_J2K50** | **11.7**<br>(273.1) | **6.0**<br>(137.3) | **8.1**<br>(0.6) | **99.9**<br>(100.0) | **41161**<br>(4586) |
| **BIN2_SUM02_J2K100** | **15.1**<br>(319.4) | **7.8**<br>(161.0) | **7.3**<br>(0.5) | **99.9**<br>(100.0) | **40963**<br>(4557) |
| **BIN2_SUM02_J2K200** | **15.7**<br>(453.1) | **8.1**<br>(227.7) | **7.4**<br>(0.3) | **99.8**<br>(99.9) | **40978**<br>(4564) |
| **BIN2_SUM02_J2K500** | **15.4**<br>(689.6) | **8.2**<br>(347.5) | **4.6**<br>(4.9) | **99.0**<br>(96.0) | **40762**<br>(4412) |
| **forced cell params / fraction 0.2** |  |  |  |  |  |
| **BIN2_SUM02_J2K1000** | **1319**<br>(667) | **691.8**<br>(369.2) | **0.1**<br>(0.1) | **98.8**<br>(97.5) | **35814**<br>(4245) |
| **forced cell params, no refine index; 0.05 indexing** |  |  |  |  |  |
| **HCOMP4** | **9.8**<br>(197) | **6.9**<br>(125.1) | **9.5**<br>(0.8) | **99.9**<br>(100.0) | **41206**<br>(4596) |
| **HCOMP8** | **11.6**<br>(423.6) | **6.0**<br>(224.2) | **8.4**<br>(0.2) | **99.8**<br>(99.9) | **41178**<br>(4598) |
| **HCOMP16** | **10.4**<br>(680.1) | **5.9**<br>(511.6) | **6.0**<br>(-0.2) | **90.8**<br>(69.2) | **37375**<br>(3422) |
| **HCOMP24** | **16.2**<br>(176.9) | **9.7**<br>(132.2) | **5.5**<br>(0.4) | **90.4**<br>(70.0) | **37159**<br>(3460) |
| **HCOMP32** | **9.3**<br>(104.1) | **5.2**<br>(102.1) | **5.5**<br>(1.0) | **66.6**<br>(14.6) | **27349**<br>(721) |
| **HCOMP64** | **14.8**<br>(0.00) | **8.4**<br>(0.00) | **3.7**<br>(-0.7) | **48.3**<br>(1.1) | **18190**<br>(52) |
| **BIN2_SUM02_HCOMP04** | **9.7**<br>(166.6) | **5.0**<br>(83.3) | **10.6** (0.9) | **99.9**<br>(100.0) | **41164**<br>(4596) |
| **BIN2_SUM02_HCOMP08** | **9.8**<br>(189.4) | **5.1**<br>(94.6) | **10.1**<br>(0.8) | **99.8**<br>(100.0) | **41169**<br>(4602) |
| **BIN2_SUM02_HCOMP16** | **12.9**<br>(516.8) | **6.6**<br>(260.0) | **7.9**<br>(0.3) | **99.8**<br>(99.7) | **40916**<br>(4549) |
| **25 % indexing** |  |  |  |  |  |
| **BIN2_SUM02_HCOMP24** | **21.2**<br>(644.9) | **10.9**<br>(323.1) | **6.3**<br>(0.2) | **99.9**<br>(99.9) | **40880**<br>(4550) |
|  | 25 % indexing |  |  |  |  |
| BIN2_ SUM02_<br>HCOMP32 | 10.4<br>(-455.6) | 5.6<br>(-244.3) | 8.0<br>(-0.2) | 96.3<br>(74.3) | 39684<br>(3426) |
|  | 25 % indexing |  |  |  |  |
| BIN2_ SUM02_<br>HCOMP64 | 9.3<br>(-169.0) | 5.2<br>(-110.1) | 6.4<br>(-0.5) | 90.0<br>(56.1) | 37101<br>(2583) |
|  | 25 % indexing |  |  |  |  |

**Table S8.** HIV reverse transcriptase structure quality metrics

| DATA | Compr. | Rwork | Rfree | Resol. | Compl. |
| --- | --- | --- | --- | --- | --- |
| Native<sup>9</sup> | 1 | 0.2136 | 0.2752 | 2.55 | 99.7 |
| CBF1<sup>10</sup> | 1 | 0.3429 | 0.4151 | 2.55 | 99.78 |
| BIN2_CBF2 | 8 | 0.3444 | 0.4109 | 2.55 | 99.69 |
| CBF40 | 40 | 0.3426 | 0.4116 | 2.55 | 92.54 |
| BIN2 | 4 | 0.3425 | 0.4124 | 2.55 | 99.7 |
| BIN4 | 16 | 0.3493 | 0.4144 | 2.55 | 99.37 |
| BIN6 | 36 | 0.4216 | 0.4025 | 2.56 | 14.9 |
| J2K200 | 100 | 0.362 | 0.4217 | 2.55 | 99.71 |
| J2K500 | 250 | 0.3645 | 0.4294 | 2.55 | 95.09 |
| J2K1000 | 504 | 0.3823 | 0.4649 | 2.55 | 67.55 |
| J2K2000 | 1008 | 0.4545 | 0.5237 | 2.55 | 72.47 |
| BIN2_CBF2_J2K50 | 200 | 0.3483 | 0.4165 | 2.55 | 99.57 |
| BIN2_CBF2_J2K200 | 798 | 0.3642 | 0.4189 | 2.55 | 96.69 |
| BIN2_CBF2_J2K500 | 2000 | 0.3969 | 0.4707 | 2.55 | 75.74 |
| HCOMP04 | 16 | 0.342 | 0.4082 | 2.55 | 99.8 |
| HCOMP08 | 107 | 0.3547 | 0.4229 | 2.55 | 99.26 |
| HCOMP16 | 850 | 0.3874 | 0.4561 | 2.55 | 87.98 |
| HCOMP24 | 985 | 0.3741 | 0.4354 | 2.55 | 86.53 |
| HCOMP32 | 1891 | 0.4006 | 0.4925 | 2.55 | 64.82 |
| HCOMP64 | 3037 | 0.3719 | 0.4929 | 2.55 | 44.15 |
| BIN2_CBF2_HCOMP04 | 33 | 0.3434 | 0.4119 | 2.55 | 99.71 |
| BIN2_CBF2_HCOMP08 | 66 | 0.3452 | 0.4145 | 2.55 | 99.69 |
| BIN2_CBF2_HCOMP16 | 253 | 0.3629 | 0.4216 | 2.55 | 94.18 |
| BIN2_CBF2_HCOMP24 | 657 | 0.356 | 0.418 | 2.55 | 93.42 |
| BIN2_CBF2_HCOMP32 | 1499 | 0.4024 | 0.4701 | 2.55 | 72.35 |
| BIN2_CBF2_HCOMP64 | 2999 | 0.4348 | 0.5082 | 2.55 | 61.2 |

<sup>9</sup> Native has the entire model in it.<sup>10</sup> CBF1 has about 30% of the model missing.

**Table S9.**
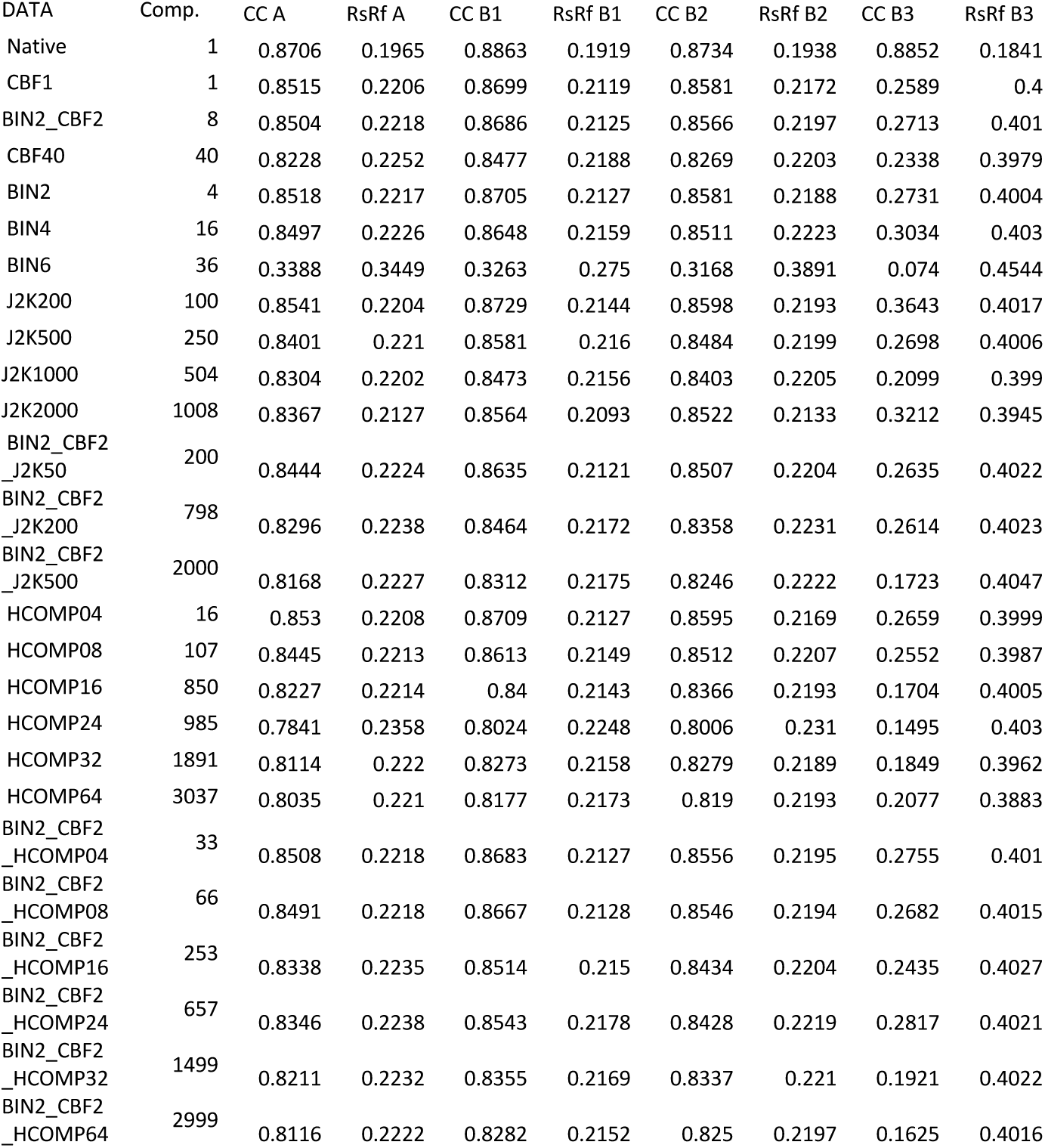
HIV reverse transcriptase map quality metrics (computed using *overlapmap*). Region A is the entire chain A in the atomic coordinate file. Regions B1, B2, and B3 are three regions in the B chain (significantly, B3 is the omit region that was not present in the phasing model). The *overlapmap* software computes the electron density difference between the comparison map (the map obtained using the compressed data) and the reference map (the map deduced from the model that was refined against the uncompressed data).

**Table S10.**
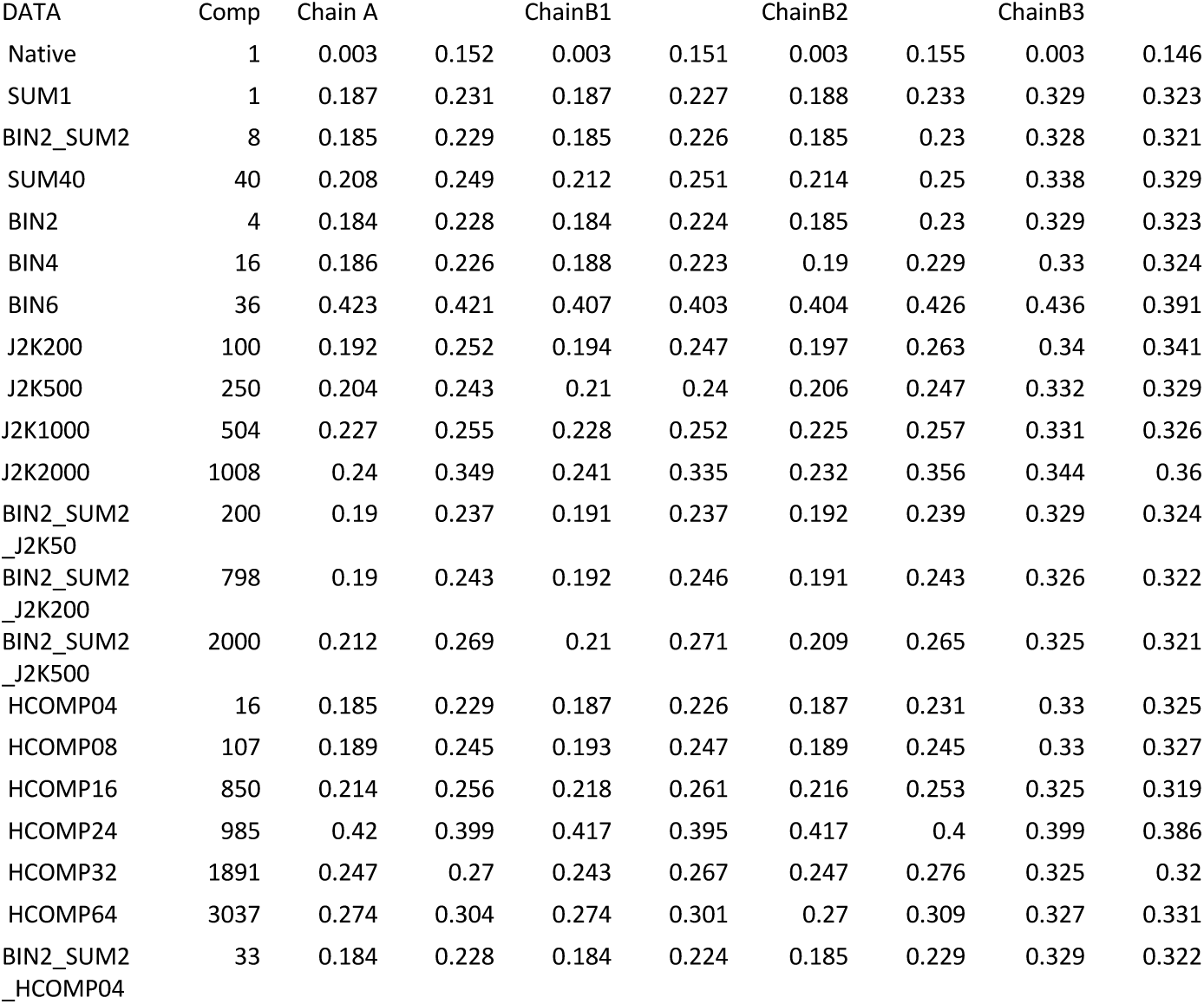
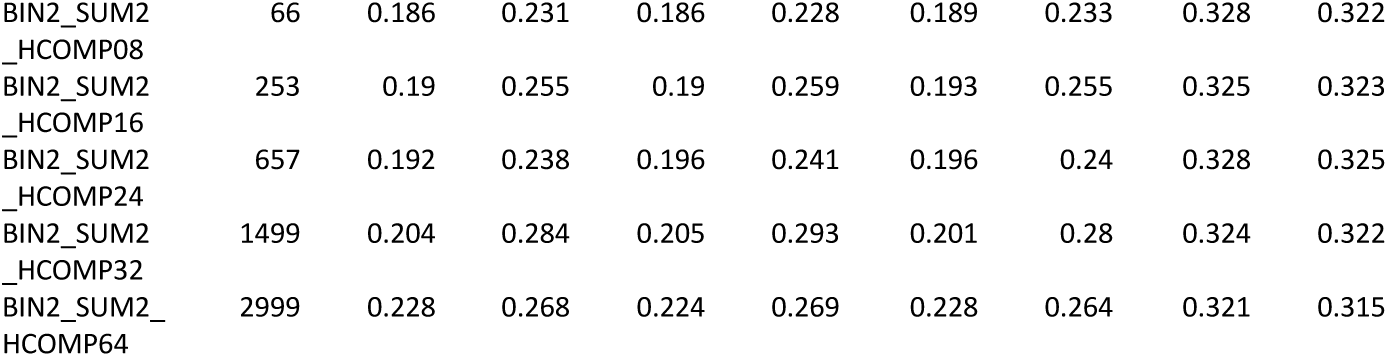
HIV reverse transcriptase map quality metrics (computed using *custom software*). Chain A is the entire chain A in the atomic coordinate file. Chains B1, B2, and B3 are three chains in the B chain (significantly, B3 is the omit region that was not present in the phasing model). In this software, the real space R value is computed in the envelope of each amino acid by adding the total differences between a comparison map and a reference map, divided by the R.M.S. variance of the reference map. Two columns are shown for each chain. In both columns, the comparison map is the map obtained using the compressed data. In the left column, the reference map is the map obtained using the uncompressed data. In the right column, the reference map is the map deduced from the model that was refined against the uncompressed data.

**Table S11.**
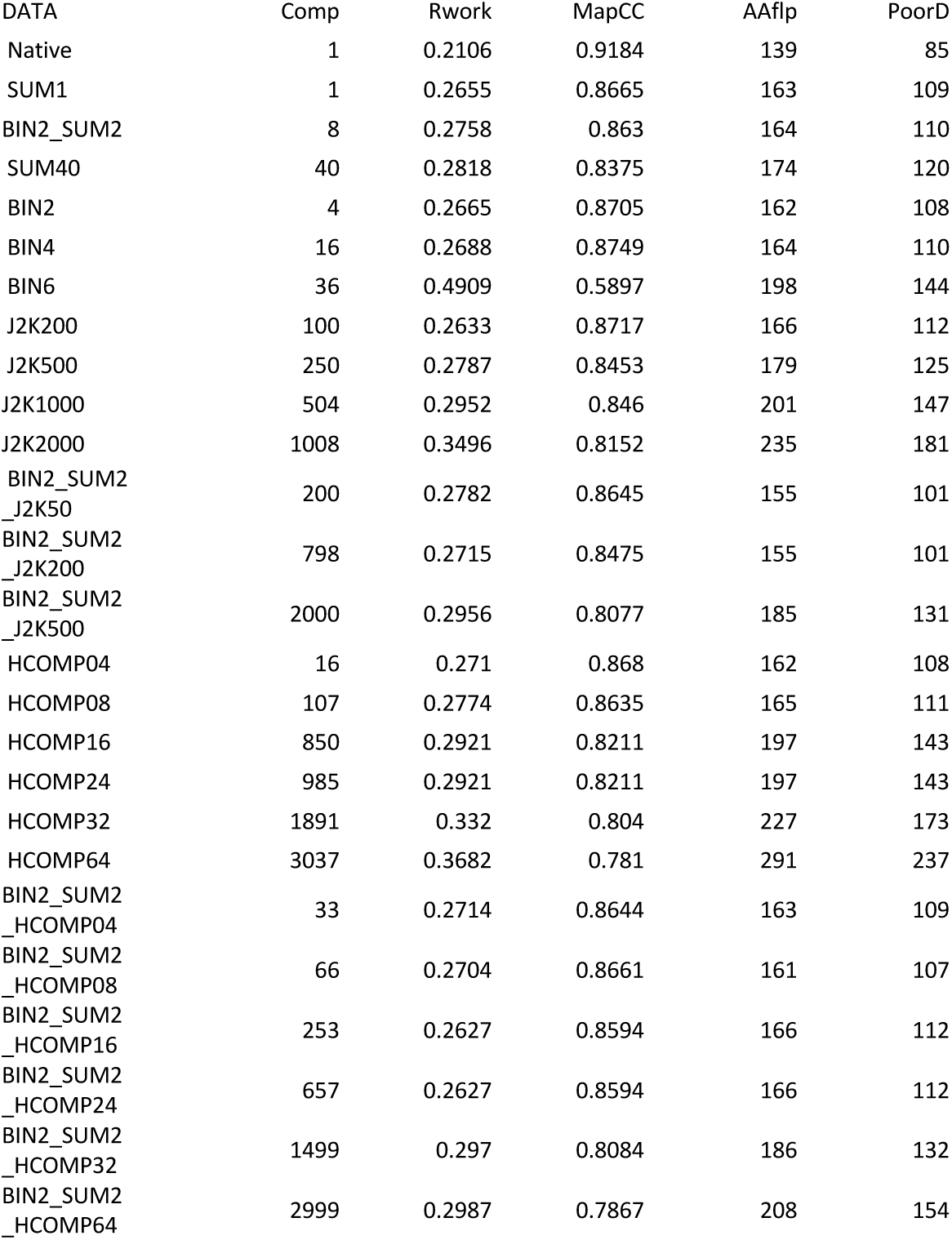
HIV reverse transcriptase map quality metrics (compared using PHENIX). MapCC is the map correlation coefficient, AAflp are the number of amino acids where the side chain may be in an incorrect conformation, and PoorD is the number of amino acids where the side chain has no conformation that has a good agreement between the model and the data.

**Figure S1.**
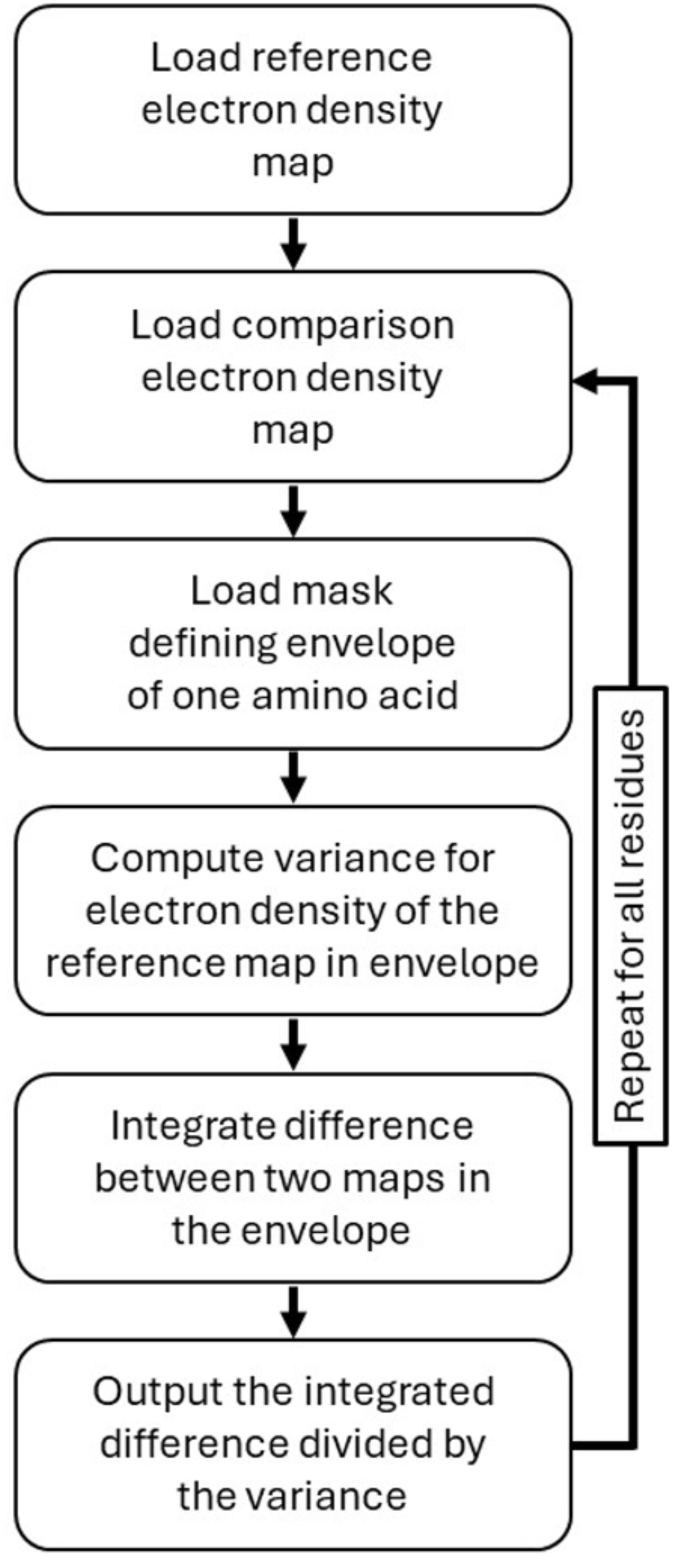
Custom software was prepared to compute real space R-values in a way that (i) permitted the comparisons of maps obtained using non-cognate atomic coordinates and (ii) permitted a custom definition of the real space R-value, such that the denominator was the variance of the reference map (rather than the total integrated electron density).

## Footnotes

1 FAIR data meet principles of Findability, Accessibility, Interoperability, and Reusability (FAIR)

2 Area Detector Systems Corp produced a popular and very successful line of Charge-Coupled-Device-based area detectors.

3 See https://bl831.als.lbl.gov/~jamesh/lossy_compression

4 Here the motive was to take as wide an image as possible (fewer films to process) while avoiding collision of spots on the film: large unit cells, smaller increments.

5 https://zenodo.org/records/12701763

6 https://github.com/dalekreitler-bnl/pa_sg_res

7 https://zenodo.org/records/12701763

8 https://github.com/nsls-ii-mx/lossy_compression_scripts

9 Native has the entire model in it.

10 CBF1 has about 30% of the model missing.

